# Haploid selection, sex ratio bias, and transitions between sex-determining systems

**DOI:** 10.1101/269431

**Authors:** Michael Francis Scott, Matthew Miles Osmond, Sarah Perin Otto

**Affiliations:** UCL Genetics Institute, Department of Genetics, Evolution and Environment, University College London, Gower Street, London WC1E 6BT; Department of Zoology, University of British Columbia, #4200 - 6270 University Boulevard, Vancouver, BC, Canada V6T 1Z4

## Abstract

Sex determination is remarkably dynamic; many taxa display shifts in the location of sex-determining loci or the evolution of entirely new sex-determining systems. Predominant theories for why we observe such transitions generally conclude that novel sex-determining systems are favoured by selection if they equalise the sex ratio or increase linkage with a locus that experiences different selection in males vs. females. We use population genetic models to extend these theories in two ways: (1) We consider the dynamics of loci very tightly linked to the ancestral sex-determining loci, e.g., within the non-recombining region of the ancestral sex chromosomes. Variation at such loci can favour the spread of new sex-determining systems in which the heterogametic sex changes (XY to ZW or ZW to XY) and the new sex-determining region is less closely linked (or even unlinked) to the locus under selection. (2) We consider selection upon haploid genotypes either during gametic competition (e.g., pollen competition) or meiosis (i.e., non-Mendelian segregation), which can cause the zygotic sex ratio to become biased. Haploid selection can drive transitions between sex-determining systems without requiring selection to act differently in diploid males vs. females. With haploid selection, we find that transitions between male and female heterogamety can evolve where linkage with the sex-determining locus is either strengthened or weakened. Furthermore, we find that sex-ratio biases may increase or decrease with the spread of new sex chromosomes, which implies that transitions between sex-determining systems cannot be simply predicted by selection to equalise the sex ratio. In fact, under many conditions, we find that transitions in sex determination are favoured equally strongly in cases where the sex ratio bias increases or decreases. Overall, our models predict that sex determination systems should be highly dynamic, particularly when haploid selection is present, consistent with the evolutionary lability of this trait in many taxa.

**Author summary:** Systems of sex determination are strikingly diverse and labile in many clades. This poses the question: what drives transitions between sex-determining systems? Here, we use models to derive conditions under which new sex-determining systems spread. Prevailing views suggest that new sex-determining systems are favoured when they equalize the sex ratio and/or when they are more closely linked to genes that experience differential selection in males and females. Our models include selection upon haploid genotypes (meiotic drive or gametic competition), which biases the sex-ratio and occurs differently in male and female gametes. Surprisingly, we find the two forces (selection to equalize the sex ratio and the benefits of hitchhiking alongside driven alleles that distort the sex ratio) will often be equally strong, and thus neither is sufficient to explain the spread of new sex-determining systems in every case. We also find that new sex-determining alleles can spread despite being less closely linked to selected loci as long as initial linkage is tight or haploid selection is present. Our models therefore predict that loci in previously unexpected genomic locations and/or experiencing various types of selection (including haploid selection) can now be implicated as drivers of transitions between sex-determining systems.

## Introduction

Animals and angiosperms exhibit extremely diverse sex-determining systems (reviewed in [1–5]). Among species with genetic sex determination (GSD), some taxa have heterogametic males (XY) and homogametic females (XX), including mammals and most dioecious plants [6]; whereas other taxa have homogametic males (ZZ) and heterogametic females (ZW), including Lepidoptera and birds. Within several taxa, the chromosome that harbours the master sex-determining locus changes, due either to translocation of the master sex-determining locus or to the evolution of a new master locus. During these transitions, the heterogametic sex can remain the same (hereafter ‘cis-GSD transitions’) as in Salmonids [7, 8], Diptera [9], and *Oryzias* [10]. Alternatively, species can switch between male and female heterogamety (XY↔ZW, hereafter ‘trans-GSD transitions’), as in snakes [11], lizards [12], eight of 26 teleost fish families [13], true fruit flies (Tephritids, [9]), amphibians [14], the angiosperm genus *Silene* [15], the angiosperm family *Salicaceae* [16, 17] and Coleoptera and Hemiptera (plate 2 [3]). Indeed, in some cases, both male and female heterogametic sex-determining systems can be found in the same species, as reported in houseflies [18], midges [19], frogs [20], cichlid fish [21], tilapia [22], sea bass [23], and lab-strains of Zebrafish [24, 25]. In addition, multiple transitions have occurred between genetic and environmental sex-determining systems (GSD↔ESD), e.g., in reptiles and fishes [5,12,13,26–29]. In sum, accumulating evidence indicates that transitions between sex-determining systems are common [4].

It has been suggested that sex-ratio selection is a particularly dominant force in the evolution of sex determination (e.g., Bull, 1983, p 66-67 [1]; Buekeboom and Perrin, 2014, Chapter 7[3]). Classic ‘Fisherian’ sex-ratio selection favours a 1:1 zygotic sex ratio when assuming that males and females are equally costly to produce [30, 31]. This follows from the fact that, for an autosomal locus, half of the genetic material is inherited from a male and half from a female [32]. Thus, if the sex ratio is biased, an individual of the rarer sex will, on average, contribute more genetic material to the next generation. Selection therefore typically favours mutants that increase investment in the rarer sex, including new sex determination systems.

The evolution of sex determination is also thought to be strongly influenced by differences in selection between the sexes [3, 33, 34]. For example, loci experiencing sexual antagonism have been shown to favour the spread of new genetic sex-determining alleles that are closely linked [35–37]. Linkage allows a stronger favourable association to build up between a male-beneficial allele and a neo-Y allele, for example. Such associations can favour cis-GSD transitions [35], trans-GSD transitions [36], and new partially-masculinizing or partially-feminizing alleles in a population with ESD [37]. By similar logic, however, existing sexually-antagonistic alleles associated with the current sex-determining locus are expected to hinder the spread of a new sex-determining system [35, 36].

One novel feature of the models developed here is that we explicitly consider the maintenance of genetic variation around the ancestral sex-determining locus (e.g., within the non-recombining region of a sex chromosome). Counterintuitively, when linkage is tight between the sex-determining locus and a selected locus, an allele good for females can be at higher frequency on the ancestral-Y than on the ancestral-X under a variety of forms of selection. In addition, selection on ancestral-X chromosomes in males can prevent the X from becoming optimally specialised for female-beneficial alleles. These factors, in turn, can favour a new ZW sex-determining locus that has weaker linkage with loci under selection, which was not revealed by previous theory [36]. A similar argument applies to ZW→XY transitions. Thus, we show that selected loci in very tight linkage with the ancestral GSD locus can favour trans-GSD transitions during which linkage associations are actually weakened.

Most significantly, we include haploid selection (gametic competition or meiotic drive) in models describing cis-GSD, trans-GSD, and GSD to ESD transitions. This poses an apparent evolutionary problem. On one hand, haploid selection is typically sex-limited in that it usually occurs among gametes produced by one sex only [38–41]. Therefore, one might expect new sex-determining systems to benefit from close linkage with haploid selected loci, as found for loci that experience diploid-sex-differences in selection [35–37]. On the other hand, associations between sex-determining loci and haploid selected loci generate biased zygotic sex ratios, which should generally hinder the spread of new sex-determining systems.

Two previous studies have considered the spread of GSD with sex-limited meiotic drive [42, 43] under a limited number of possible genetic architectures and diploid selective regimes. Ubeda et al. (2015) [43] considered ancestral-ESD (with no sex-ratio bias) and numerically showed that new GSD alleles can spread if they arise in linkage with meiotic drive loci. For example, a masculinizing allele spreads in association with an allele that is favoured during male meiosis, causing sex ratios to become male-biased. This suggests that the benefits of associating with driving alleles can overwhelm selection to balance the sex ratio. However, Kozielska et al. (2010) [42] considered an ancestral GSD system that is perfectly linked to a meiotic driver and therefore exhibiting an ancestral sex ratio bias. They found that a new, completely unlinked, GSD system can spread if it generates the rarer sex, creating a balanced sex ratio. This suggests that Fisherian sex-ratio selection can overwhelm the benefits of being associated with driving alleles. It is thus currently unclear when haploid selection favors increased versus decreased linkage between haploid selected loci and a new sex-determining locus. In addition, because the sex ratio is determined by linkage between haploid selected loci and the sex-determining locus, it is also unclear when Fisherian sex-ratio selection is the most important driver of transitions between sex-determining systems.

Here, we analytically find the conditions under which new GSD or ESD systems spread in ancestral GSD systems with any degree of linkage between the loci involved and arbitrary forms of haploid and diploid selection. Doing so, we reconcile and generalize the results of Kozielska et al. (2010) [42] and Ubeda et al. (2015) [43] by deriving conditions for the spread of new GSD systems that alter linkage with haploid selected loci. Our result is qualitatively distinct from those for diploid selection alone [35, 36] and suggests that haploid selection is more likely to promote transitions between sex-determination systems. We also show that transitions involving haploid selection cannot be simply explained by invoking sex-ratio selection. In particular, under a wide range of conditions, we show that transitions in sex-determining system are favoured *equally strongly* in situations where sex-ratio biases increase or decrease (and in situations where sex-ratio biases are ancestrally present or absent). Finally, we show that ESD may not evolve, even if the sex ratio is initially biased by haploid selection, which is not predicted by previous theories for transitions to ESD [1, 31, 32]. Together, our results suggest that both selection to equalize the sex ratio and the benefits of associating with haploid selected alleles can drive transitions between sex-determining systems, leading to stronger or weaker sex-linkage and increased or decreased sex ratio bias.

## Model

We consider transitions between ancestral and novel sex-determining systems using a three-locus model, each locus having two alleles (Fig 1). A full description of our model, including recursion equations, is given in S1 Appendix. Locus **X** is the ancestral sex-determining region, with alleles X and Y (or Z and W). Locus **A** is a locus under selection, with alleles *A* and *a*. Locus **M** is a novel sex-determining region, at which the null allele (*M*) is initially fixed in the population such that sex of zygotes is determined by the genotype at the ancestral sex-determining region, **X**; XX genotypes become females and XY become males (or ZW become females and ZZ become males). To evaluate the evolution of new sex-determining systems, we consider the spread of a novel sex-determining allele (*m*) at the **M** locus.

**Fig 1.**
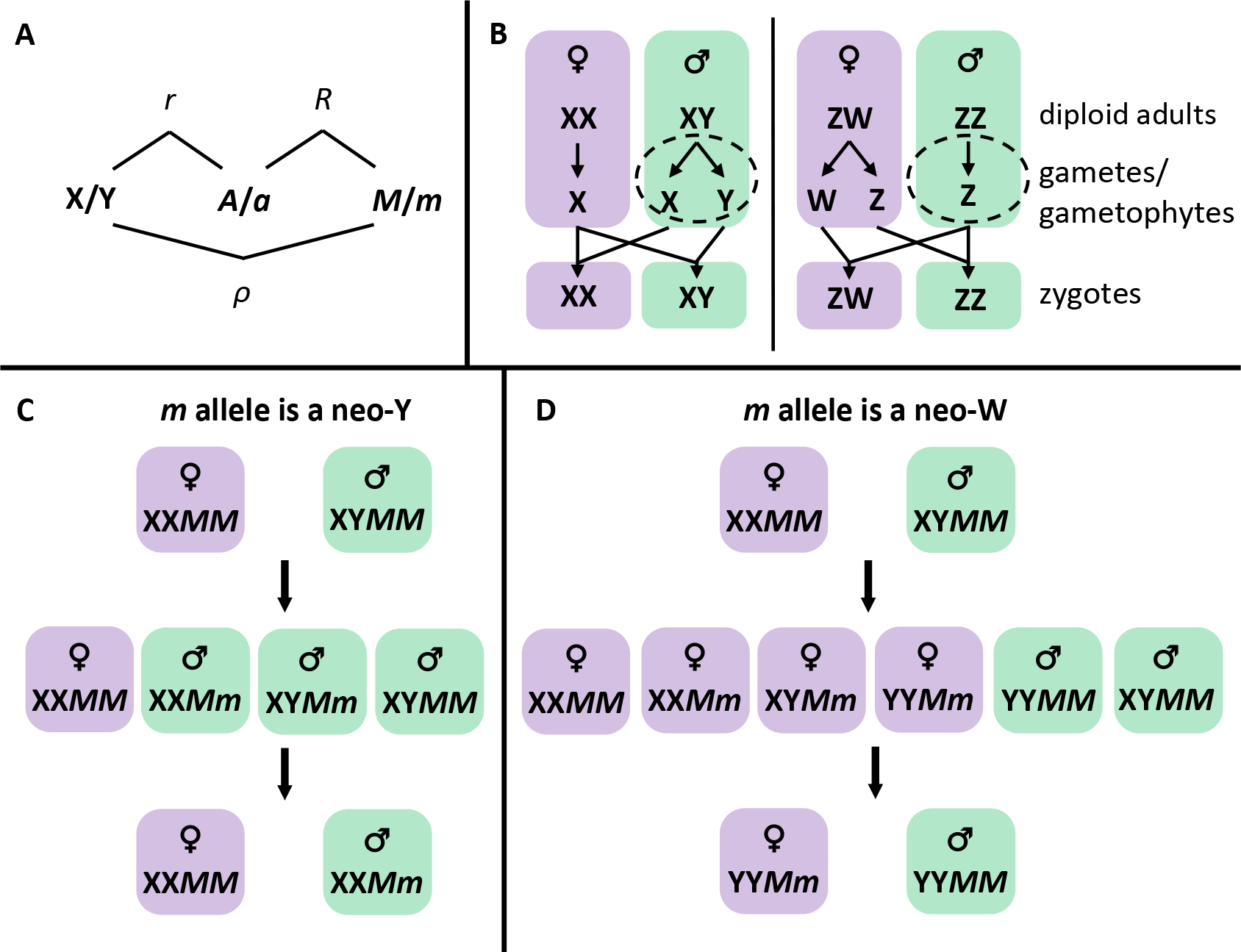
Outline of model features. Panel A: Recombination rate parameters between the ancestral-sex-determining locus (**X**, here assumed to have alleles X and Y), a locus under selection (**A**, with alleles *A* and *a*), and a new sex-determining locus (**M**, with alleles *M* and *m*). Panel B: Haploid selection is often sex-limited, occurring during haploid production or competition in one sex (shown here in males by dashed circles). If X or Y alleles are linked with alleles that experience haploid selection in males (*r* < 1 /2), then zygotic sex ratios can become biased because either X- or Y-bearing male gametes/gametophytes will be more abundant after haploid selection. Similarly, zygotic sex-ratio biases can arise if haploid selected alleles are linked with new sex-determining alleles (*R* < 1 /2). However, the zygotic sex ratio is not biased by male haploid selection in ZW sex-determining systems. Panel C: During cis-GSD transitions (XY to XY or ZW to ZW), a neo-Y allele (*m*) spreads to pseudo-fixation (i.e, all males bear the neo-Y) and the ancestral Y allele is lost. Panel D: During trans-GSD transitions (XY to ZW or ZW to XY), a neo-W allele (*m*) spreads to pseudo-fixation (i.e, all females bear the neo-W) and the ancestral X allele is lost. Neo-W alleles allow Y-associated alleles into females, which may impede or aid their spread.

Here, we assume that the **M** locus is ‘epistatically dominant’ over the **X** locus such that zygotes with at least one *m* allele develop as females with probability *k* and as males with probability 1 - *k*, regardless of the **X** locus genotype. With *k* = 0, the *m* allele is a masculinizer (a neo-Y allele) and with *k* =1 the *m* allele is a feminizer (a neo-W allele). With intermediate *k*, we can interpret *m* as an environmental sex-determination (ESD) allele, such that zygotes develop as females in a proportion (*k*) of the environments they experience. The assumption that derived sex-determining loci are epistatically dominant is motivated by empirical systems in which multiple sex determining alleles segregate (i.e., X, Y, Z, and W alleles present), such as, cichlid fish [21], platyfish (*Xiphophorus maculatus* [44]), houseflies (*Musca domestica* [45]), western clawed frogs (*Xenopus tropicalis* [46]) and *Rana rugosa* [20]. Nevertheless, our supplementary analysis file (S1 File) allows other dominance relationships between loci to be specified (see also [35] supplementary material for a numerical analysis).

We consider two forms of selection upon haploid genotypes, ‘gametic competition’ and ‘meiotic drive’. During gametic competition, we assume that a representative sample of all gametes/gametophytes (hereafter gametes) compete with others of the same sex for fertilization, which implies a polygamous mating system. Relative fitnesses in sex ∘ ∈ {♀, ♂} during gametic competition are given by 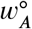 and 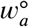 (see Table 1). On the other hand, meiotic drive in our model only affects the segregation of gametes produced by heterozgotes. Specifically, gametes produced by *Aa* heterozgotes of sex ∘ bear allele *A* with probability *α*^∘^. We note that competition between sperm produced by a single male (e.g., in a monogamous mating system) would be appropriately modelled as male meiotic drive, as only the frequency of gametes produced by heterozygotes would be affected. However, we do not consider scenarios in which there is competition among gametes produced by a small number of males/females (e.g., [47]).

**Table 1.** **Relative fitness of different genotypes in sex,** ° ∈ {♀, ♂}

| Genotype | Relative fitness during gametic competition |
| --- | --- |
| $A$ | $w_A^\circ = 1 + t^\circ$ |
| $a$ | $w_a^\circ = 1$ |
| Genotype | Relative fitness during diploid selection |
| $AA$ | $w_{AA}^\circ = 1 + s^\circ$ |
| $Aa$ | $w_{Aa}^\circ = 1 + h^\circ s^\circ$ |
| $aa$ | $w_{aa}^\circ = 1$ |
| Genotype | Transmission during meiosis in $Aa$ heterozygotes |
| $A$ | $\alpha^\circ = 1/2 + \alpha_\Delta^\circ/2$ |
| $a$ | $1 - \alpha^\circ = 1/2 - \alpha_\Delta^\circ/2$ |

In each generation, we census the genotype frequencies in male and female gametes before gametic competition. After gametic competition, conjugation between male and female gametes occurs at random. The resulting zygotes develop as males or females, depending on their genotypes at the **X** and **M** loci. Diploid males and females then experience viability and/or individual-based fertility selection, with relative fitnesses 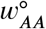, 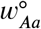, and 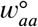. We do not consider fertility selection that depends on the mating partner, e.g., sexual selection with variation in choosiness. The next generation of gametes is produced by meiosis, during which recombination and sex-specific meiotic drive can occur. Recombination (i.e., an odd number of cross-overs) occurs between loci **X** and **A** with probability *r*, between loci **A** and **M** with probability *R*, and between loci **X** and **M** with probability *ρ*. Any linear order of the loci can be modelled with appropriate choices of *r*, *R*, and *ρ* (see Fig 1A and Table S1). Our model is entirely deterministic and hence ignores chance fluctuations in allele frequencies due to genetic drift.

The model outlined above describes both ancestral XY and ZW sex-determining systems. Without loss of generality, we refer to the ancestrally heterogametic sex as male and the ancestrally homogametic sex as female. That is, we primarily describe an ancestral XY sex-determining system but our model is equally applicable to an ancestral ZW sex-determining system (relabelling the ancestrally heterogametic sex as female and the ancestrally homogametic sex as male and switching the labels of males and females throughout). We use a superscript to specify the ancestral sex-determining system described, e.g.,^(XY)^ for ancestral XY sex-determination.

In the ancestral population, it is convenient to follow the frequency of the *A* allele among female gametes (eggs), 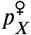, and among X-bearing, 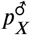, or among Y-bearing, 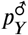, male gametes (sperm/pollen). We also track the fraction of male gametes that are Y-bearing, *q*, which may deviate from 1/2 due to meiotic drive in males. We consider only equilibrium frequencies of alleles, 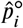, and Y-bearing male gametes, 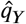, when determining the invasion of new sex-determining factors. We use *ζ* to measure the sex ratio (fraction male) among zygotes, which is determined by the allele frequencies and haploid selection coefficients (Table S2).

## Results

We begin by describing the general conditions under which new genetic sex determining alleles can spread within a population, without explicitly specifying ancestral allele frequencies. These general conditions then allow us to consider several special cases of interest in subsequent sections, where equilibrium ancestral allele frequencies are explicitly calculated. Finally, we consider the spread of alleles that specify environmental sex determination.

### Generic invasion by a neo-Y or neo-W

The evolution of a new sex-determining system requires that a rare mutant allele, *m*, at the novel sex-determining locus, **M**, increases in frequency when rare. Specifically, *m* invades when 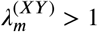, where 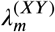 is the leading eigenvalue of the system of eight equations describing *m*-bearing gamete frequencies, S1.1. This system simplifies substantially for an epistatically dominant neo-Y (*k* = 0) or neo-W (*k* = 1), see S3 Appendix for details.

Invasion by a neo-Y or a neo-W primarily depends on the “haplotypic growth rates” (denoted by 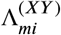) of the neo-sex determination allele *m* on background *i* ∈ {*A, a*} without accounting for loss due to recombination (*R* = 0), see Table 2. If both haplotypic growth rates are greater than one 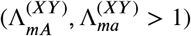, then the new sex-determining allele invades regardless of the rate of recombination between the new sex-determining locus and the selected locus (*R*). Conversely, if both haplotypic growth rates are less than one 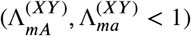, then invasion can never occur. Finally, if only one haplotypic growth rate is greater than one, the new sex-determining allele can always invade when arising at a locus that is tightly linked to the selected locus (*R* ≈ 0). Furthermore, it can be shown that the leading eigenvalue declines with recombination rate, *R*, and invasion requires that *R* is sufficiently small such that:

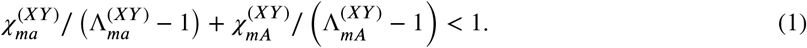

**Table 2.**
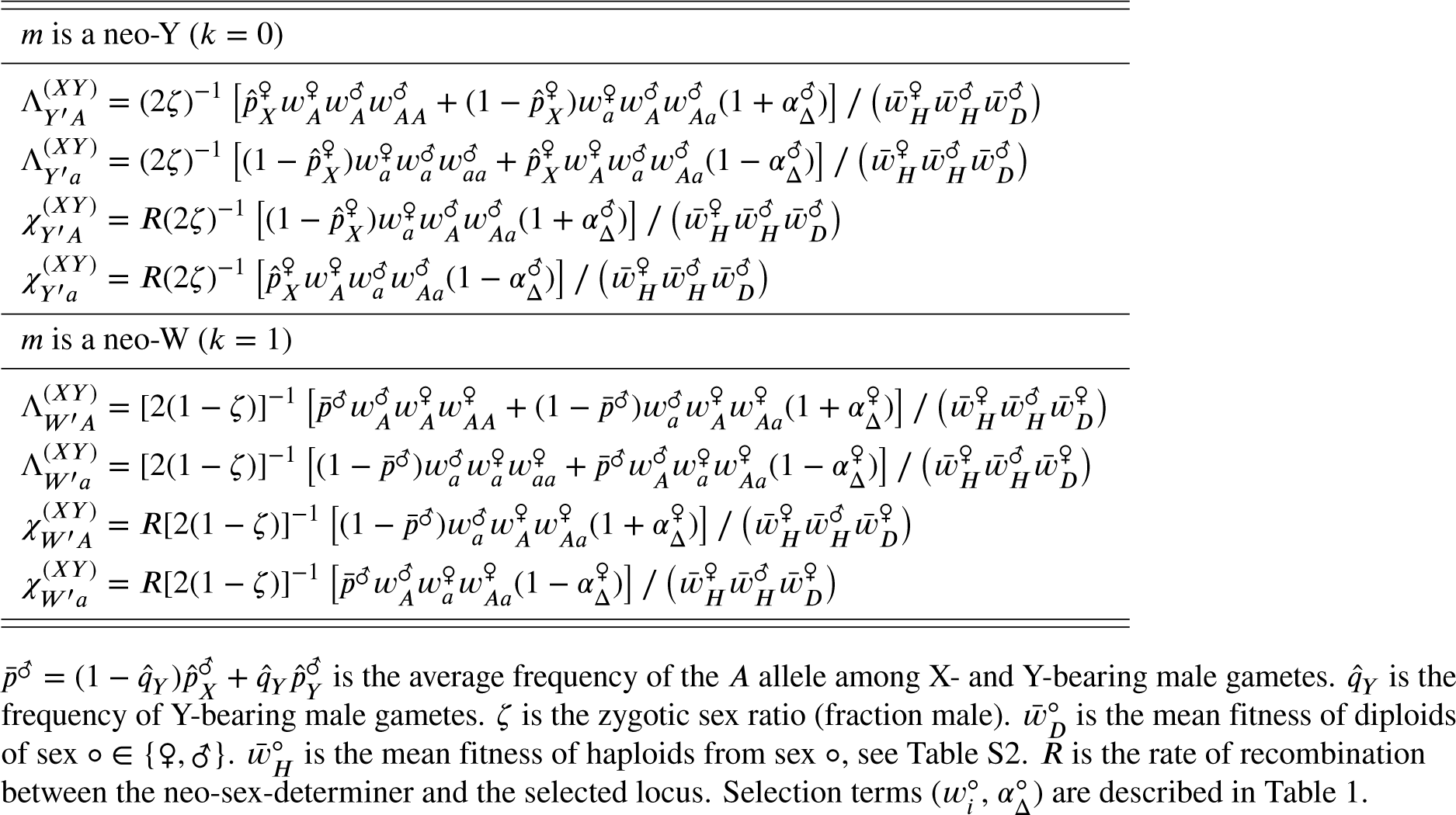
Parameters determining invasion of mutant neo-Y and neo-W alleles into an ancestrally XY system

Here 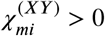 is the rate at which mutant haplotypes on background *i* ∈ {*A, a*} recombine onto the other **A** locus background in heterozygotes (which is proportional to *R*, see Table 2). This is a “dissociative force” that breaks down linkage disequilibrium.

Condition 1 may or may not be satisfied for the full range of locations of the new sex-determining locus, including *R* =1/2 (e.g., on an autosome), depending on the nature of selection. Interpreting this condition, if we assume that only the *mA* haplotype would increase in frequency when *R* = 0 (i.e., 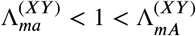), then the first term on the left-hand side of (1) is negative and invasion requires that growth rate of *mA* haplotypes 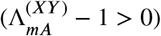 and the rate at which they are produced by recombination 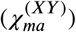 are sufficiently large relative to the rate of decline of *ma* haplotypes 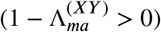 and the rate at which *m* and *A* are dissociated by recombination 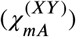.

The haplotypic growth rates and dissociative forces are listed in Table 2 for a neo-Y and neo-W invading an ancestrally XY system. From this table and the arguments above we draw four main points about the generic invasion of neo-Y and neo-W mutations (without specifying the ancestral equilibrium): (1) Fisherian sex-ratio selection will favour the spread of a neo-W and inhibit the spread of a neo-Y if the ancestral zygotic sex ratio is biased towards males (i.e., the first factor of the 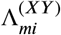 is greater than one for a neo-W and less than one for a neo-Y when *ζ* > 1/2). Thus, neo-sex-determining alleles that specify the rarer sex are favoured by Fisherian sex ratio selection. (2) In addition, the new sex determining allele has associations with alleles favored by either haploid or diploid selection (terms in square brackets). Importantly, invasion by a neo-Y (neo-W) does not directly depend on the fitness of female (male) diploids. This is because a dominant neo-Y (neo-W) is always found in males (females), and therefore the frequency of the neo-Y (neo-W), *m*, only changes in males (females), Fig 1C,D. (3) Haploid selection thus plays two roles, generating Fisherian selection to equalize the ancestral sex-ratio (through *ζ*) and generating selection for the neo-Y/neo-W through associations with haploid selected loci, which can distort the sex ratio. Each role influences the invasion dynamics of a new sex-determining allele, allowing the sex ratio to become more or less biased during a transition (as previously found in two special cases; [42, 43]). (4) Finally, Table 2 shows that the genetic contexts that arise during cis- and trans-GSD transitions are qualitatively different. This is because, in an ancestrally XY system, a gamete with the neo-Y always pairs with a female gamete containing an X, Fig 1C. By contrast, a gamete with a neo-W can pair with an X- or Y-bearing male gamete, Fig 1D. Consequently, neo-W-bearing females obtain a different frequency of *A* alleles from mating compared to ancestral (*MM*) females (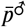 versus 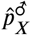, respectively). This can inhibit or favour the spread of a neo-W.

In order to explicitly determine the conditions under which a new sex-determining allele spreads, we next calculate the equilibrium frequency of the *A* allele (i.e., 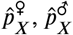, and 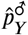) and Y-bearing male gametes 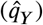 in the ancestral population. Because only the **A** locus experiences selection directly, any deterministic evolution requires that there be a polymorphism at the **A** locus. Polymorphisms can be maintained by mutation-selection balance or occur transiently during the spread of beneficial alleles. Here, however, we focus on polymorphisms maintained by selection for longer periods. Such polymorphisms can be maintained by heterozygote advantage, sexually-antagonistic selection, ploidally-antagonistic selection, or a combination [48]. We analytically calculate equilibrium frequencies using two alternative simplifying assumptions: (1) the **A** locus is tightly linked to the non-recombining region around the ancestral sex-determining locus (*r* ≈ 0) or (2) selection is weak relative to recombination (*s*∘, *t*∘, 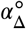 ≪ *r*). The ancestral equilibria and their stability conditions are given in S2 Appendix.

### Tight linkage with the ancestral sex-determining locus (*r* ≈ 0)

When there is complete linkage between the ancestral sex-determining locus and the selected locus **A** (*r* = 0), either the *A* allele or the *a* allele must be fixed in gametes containing a Y allele (S2 Appendix). Because the labelling of alleles is arbitrary, we will assume that the *a* locus is fixed in gametes with a Y 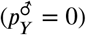, without loss of generality. If there are two alleles maintained at the **A** locus, the *A* allele can be fixed 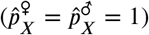 or segregating at an intermediate frequency 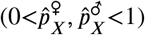 in gametes with an X.

We find that a neo-Y allele can never invade an ancestral XY system that already has tight linkage with the locus under selection (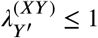 when *r* = 0; for details see S1 File). In essence, through tight linkage with the **A** locus, the ancestral Y becomes strongly specialized on the allele that has the highest fitness across male haploid and diploid phases. It is thus not possible for a neo-Y to create males that have higher fitness than the ancestral Y, and cis-GSD transitions are never favoured.

Neo-W alleles, on the other hand, can invade an ancestral XY system (the full invasion conditions are given in S3 Appendix; equations S3.1 and S3.2). Invasion occurs when neo-W females can have higher fitness than the XX females in the ancestral population. Neo-W invasion is possible under all forms of selection that can maintain a polymorphism (sexually-antagonistic selection, overdominance, ploidally-antagonistic selection, or some combination, e.g., Fig S2, Fig S3, and Fig S8). Thus,

*Conclusion 1:* **Selection on loci in or near the non-recombining region around the ancestral sex-determining locus (r ≈ 0) prevents cis-GSD transitions (XY ↔ XY, ZW ↔ ZW) but can spur trans-GSD transitions (XY ↔ ZW).** To clarify conditions under which trans-GSD transitions can occur, we focus here on cases where there is no haploid selection (and hence no sex-ratio bias) and discuss the additional effect of haploid selection in S3 Appendix. Broadly, it is possible for neo-W females to have higher fitness than XX females for two reasons. Firstly, because the ancestral X experiences selection in both males and females, the X may be unable to specialize strongly on an allele favoured in females. Secondly, an allele can be associated with the Y and yet favoured in females. In turn, a neo-W can spread because (a) it is only found in females and is therefore unleashed from counterselection in males (corresponding to 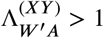), (b) it allows Y-associated alleles into females (corresponding to 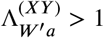).

We first give an example where neo-W-A haplotypes can spread because the neo-W is unleashed from counterselection in males (case (a), where 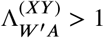). When *A* is female beneficial and *a* is male beneficial, the *A* allele can be fixed 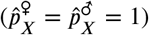 or polymorphic 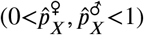 on the X. In this case, polymorphism on the ancestral-X indicates suboptimal specialisation for females fitness, which occurs because the *A* allele is counterselected in males (requires that 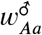 sufficiently small relative to 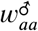). Neo-Ws, however, spend no time in males and can build stronger associations with the female-beneficial *A* allele, allowing them to spread (see gray region in Fig 2A).

**Fig 2.**
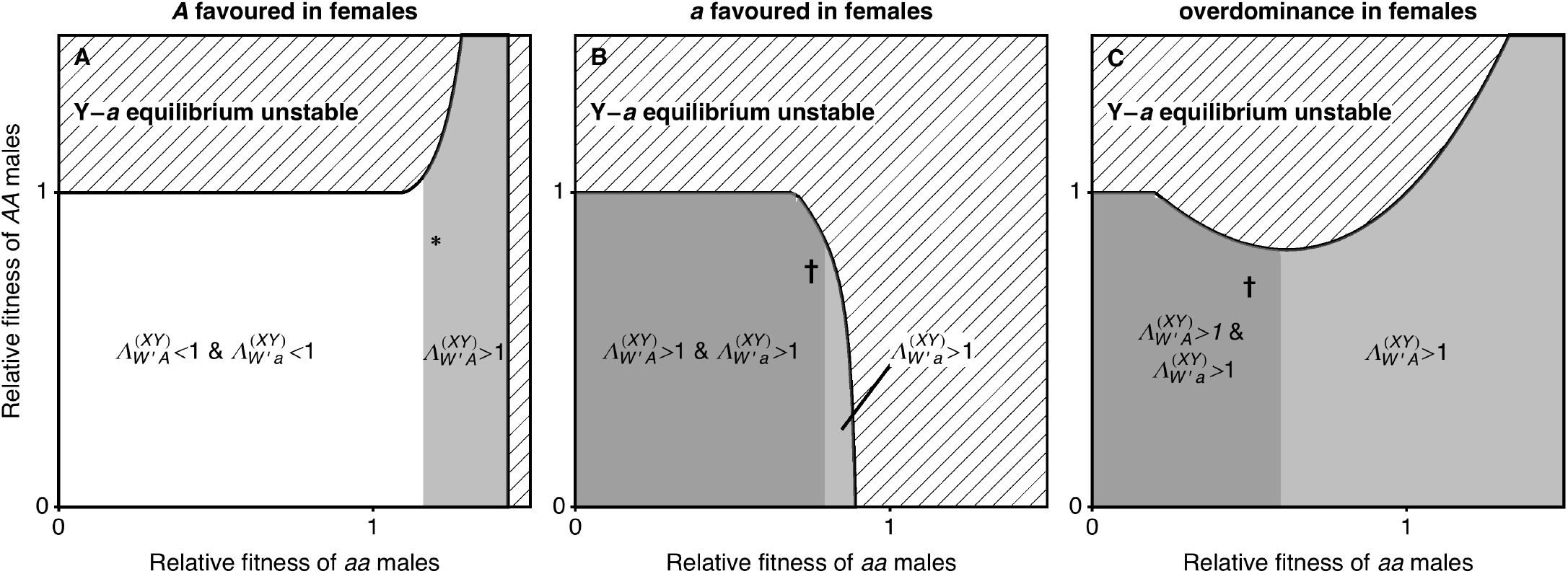
When the ancestral XY locus is tightly linked to a locus under selection (*r* = 0), one or both neo-W haplotypes can spread (no haploid selection). We vary the fitness of male homozygotes relative to heterozygotes 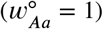 and only consider stable equilibria at which both **A** locus alleles are maintained and the *a* allele is initially fixed on the Y (non-hatched region). Here, selection in females can favour the *A* allele (panel A, 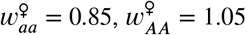), favour the *a* allele (panel B, 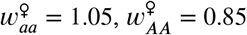), or be overdominant (panel C, 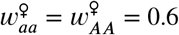). If either haplotypic growth rate (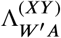 or 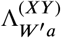) is greater than one, then a rare neo-W allele can spread for, at least, some values of *R* > *r* (grey regions). The parameter values marked with an asterisk correspond to the fitnesses used in Fig 3C. Fig S1 shows the dynamics arising with the parameters marked with a dagger.

**Fig 3.**
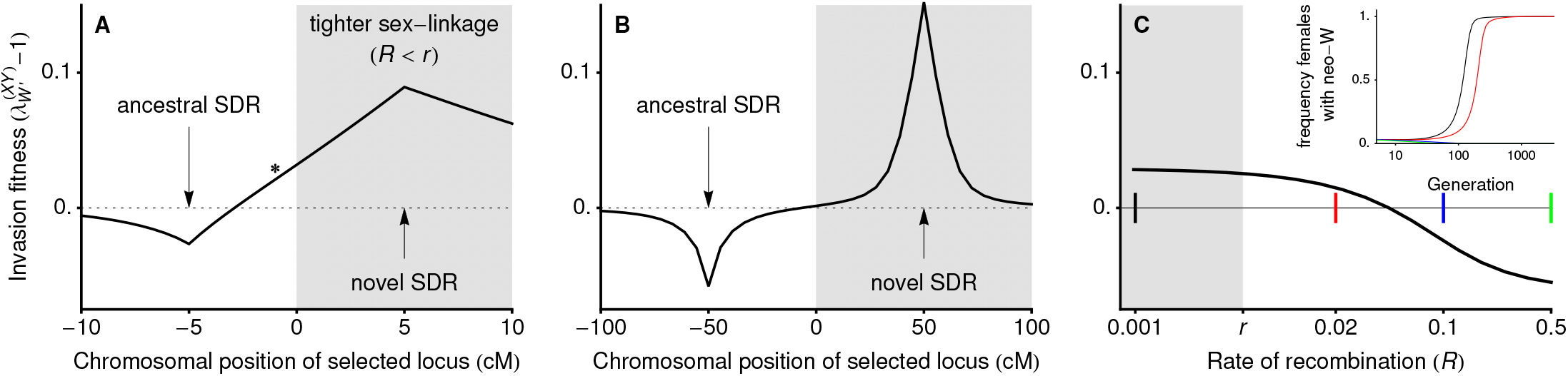
Transitions between XY and ZW systems can occur even when the new sex-determining locus is less tightly linked to a locus under sexually-antagonistic selection (no haploid selection). In panel A, linkage is initially tight relative to selection and a neo-W can invade even when it is less tightly linked with the selected locus (*r* < *R*; unshaded region around *). In panel B, linkage is loose enough relative to selection that the analytical results assuming weak selection hold, and a neo-W allele can only invade when it arises at a locus more tightly linked with the selected locus (*R* < *r*; shaded region). In panel C we vary the recombination rate between the neo-W and the selected locus (*R*) for a fixed recombination rate between the ancestral sex-determining locus and the selected locus (r = 0.005). Coloured markers show recombination rates for which the temporal dynamics of invasion are plotted in the inset (frequency of females carrying a neo-W), demonstrating that neo-W alleles can reach fixation if they are more (black) or less (red) closely linked to a locus experiencing sexually-antagonistic selection. A very loosely linked neo-W does not spread in this case (blue and green lines overlap and go to 0 in inset). Fitness parameters are: 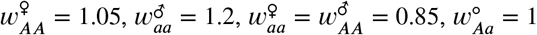.

We next give an example where neo-W-*a* haplotypes can spread because they bring in female beneficial alleles associated with the Y (case (b), where 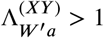). When there is overdominance in males, X-*A* Y-*a* males have high fitness and the *A* allele is favoured by selection on the X background in males. Therefore, the *A* allele can be polymorphic or even fixed on the X background despite selection favouring the *a* allele in females (e.g., see non-hatched region in Fig 2B and [49, 50]). In such cases, neo-W-*a* haplotypes can spread because they create more *Aa* and *aa* females when pairing with an X-bearing gamete from males and because they bring more of the Y-*a* haplotype into females, where it has higher fitness (Fig 1D).

In some cases, both neo-W-*A* and neo-W-*a* haplotypes can spread. For example, when *AA* individuals have low fitness in females yet the *A* is polymorphic or fixed on the X background due to overdominance in males (Fig 2B and 2C), both neo-W-*A* and neo-W-*a* haplotypes produce fewer unfit *AA* females. This is true for the neo-W-*A* haplotype because it can pair with a Y-*a* haplotype and still be female. Whenever both haplotypic growth rates are greater than one, invasion by a neo-W is expected regardless of its linkage with the selected locus (i.e., for any *R*), see Fig S1 and Fig S2 for examples. As a consequence, evolution can favor a new sex determination system on a different chromosome (*R* =1/2), despite the fact that this unlinks the sex-determining locus from the selected locus.

When only one neo-W haplotype has growth rate greater than one (see Fig 2), a neo-W allele can invade as long as Eq (1) is satisfied, which may require that the recombination rate, *R*, is small enough. Nevertheless, because we assume here that *r* is small, these results indicate that a more loosely linked sex-determining region (*r* < *R*) can spread. For example, tightly sex-linked loci that experience sexually-antagonistic selection can drive trans-GSD transitions in which the new sex-determining locus is less closely linked (*R* > *r*, Fig 3), but the analysis in S1 File indicates that a new unlinked sex-determining allele (*R* = 1/2) cannot invade when selection is purely sexually-antognistic (directional selection in each sex and no haploid selection).

Assuming selection is weak relative to recombination, van Doorn and Kirkpatrick [36] showed that invasion by a neo-W allele occurs under the same conditions as its fixation in females. An equivalent analysis is not possible where recombination rates are low. However, numerical simulations demonstrate that, with tight sex linkage, neo-Y or neo-W alleles do not necessarily reach fixation in males or females, respectively, which can lead to the stable maintenance of a mixed sex-determining system, in which X, Y, and neo-W alleles all segregate (e.g., Fig S9B,C).

From the arguments above we reach:

*Conclusion 2:* **With tight linkage between a selected locus and the ancestral sex-determining locus (*r* ≈ 0), trans-GSD transitions (XY ↔ ZW) can be favoured by selection even if they weaken sex-linkage (*r* < *R*), potentially shifting sex determination to a different chromosome (*R* =1/2).** Such transitions can also lead to the maintenance of multifactorial sex-determination systems. With haploid selection, Conclusions 1 & 2 continue to apply (S3 Appendix). The parameters for which neo-W-A and neo-W-a haplotypes spread under various forms of haploid selection are plotted in Fig S4, Fig S5, Fig S6, Fig S7. In particular, we note that adding haploid selection allows shifts in sex determination to a different chromosome (*R* = 1/2) even when selection is sexually antagonistic with directional selection in each diploid sex, e.g., Fig S3. Furthermore, haploid selection allows variation to be maintained by ploidally-antagonistic selection, under which trans-GSD transitions may also be favoured, Fig S8. Some cases of XY → ZW transitions where *r* = 0, *R* = 1/2, and selection is ploidally-antagonistic (meiotic drive in males opposed by diploid selection) were studied by Kozielska et al. [42], who found that sex-ratio biases are reduced during these transitions. However, such transitions are not always driven by selection to reduce sex-ratio bias. For example, with XY sex determination and haploid selection in females, sex ratios are not ancestrally biased yet a neo-W can invade (Fig S8). We further discuss how the spread of neo-sex-determining alleles is influenced by associations with haploid selected loci in the next section.

### Loose linkage with the ancestral sex-determining region

Here we assume that selection is weak (*s*^∘^, *t*^∘^, 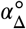 of order *ϵ*, where *ϵ* is some number much less than one) and thus implicitly assume that all recombination rates (*r*, *R* and *ρ*) are large relative to selection. To leading order in selection,

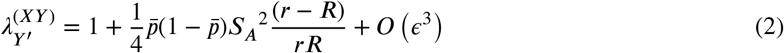

and

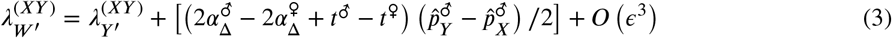

where 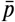 is the frequency of A, to leading-order (Eq S2.3), and 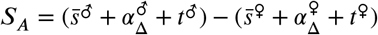 describes sex differences in selection for the *A* versus *a* allele across diploid selection, meiosis, and gametic competition. The diploid selection term, 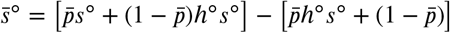, is the difference in fitness between *A* and *a* alleles in diploids of sex ∘ ∈ {♀, ♂ }. The difference in *A*-allele-frequency among Y-bearing sperm versus X-bearing sperm is, at equilibrium, 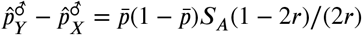.

Eq (2) demonstrates that, under weak selection, a neo-Y allele will invade an XY system 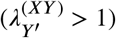 if and only if it is more closely linked to the selected locus than the ancestral sex-determining locus (i.e., if *R* < *r*). This echoes our results above where a neo-Y could never invade if *r* ≈ 0. It is also consistent with the results of [35], who considered diploid selection only and also found that cis-GSD transitions can only occur when the new sex-determining locus is more closely linked to a locus under sexually-antagonistic selection.

*Conclusion 3A:* **New sex-determining alleles (causing cis-GSD transitions, XY ↔ XY or ZW ↔ ZW) are favoured if they arise more closely linked with a locus that experiences (haploid and/or diploid) selection than the ancestral-sex-determining locus (*R* < *r*).** Similarly, in the absence of haploid selection 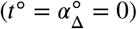, Eq (3) indicates that trans-GSD transitions can occur if and only if the new sex-determining locus is more closely linked to a locus under selection, *R* < *r*, as found by [36]. With haploid selection, a neo-W is also usually favoured when it is more closely linked to the selected locus than the ancestral sex-determining region is, (*R* < *r*, e.g., Figs 3B and 4); this is true unless the last term in Eq (3) is negative and dominant over the first, which requires relatively restrictive combinations of selection and recombination parameters. For example, with haploid selection, a neo-W will always be favoured if it arises in linkage with a selected locus (*R* < 1/2) that is ancestrally autosomal (*r* = 1/2, leading to 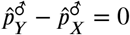).

**Fig 4.**
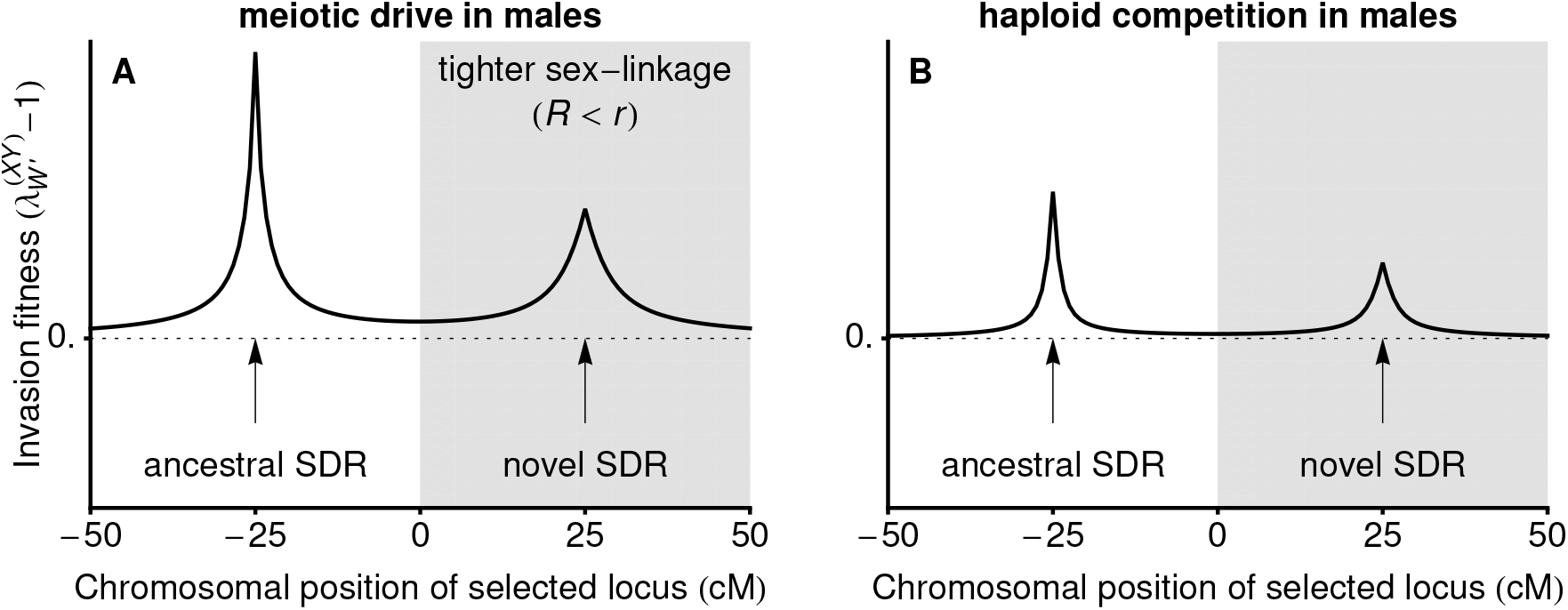
Ploidally-antagonistic selection allows a less tightly linked neo-W allele to invade. In panel A, male drive 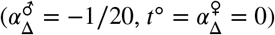 opposes selection in diploids (no sex-differences: *s*^∘^ = 1/10, *h*^∘^ = 7/10). In panel B, gametic competition in males (*t*^♂^ = −1/10, 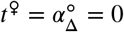) opposes selection in diploids (sex-differences: *s*^♂^ = 3/20, *s*^♀^ = 1/20, *h*^∘^ = 7/10). In either case the new sex-determining allele can invade regardless of *R*, even when linkage to the selected locus is reduced (white regions).

*Conclusion 3B:* **New sex-determining alleles (causing trans-GSD transitions, XY ↔ ZW) are usually favoured if they arise more closely linked with a locus that experiences (haploid and/or diploid) selection than the ancestral-sex-determining locus (*R* < *r*).**

However, with haploid selection and some ancestral sex-linkage (*r* < 1/2; allowing allele frequency differences on the X and Y), the term in square brackets in Eq (3) can be positive. This leads to

*Conclusion 3C:* **With haploid selection, new sex-determining alleles (causing trans-GSD transitions, XY ↔ ZW) can spread even if they arise less closely linked with a locus that experiences selection than the ancestral-sex-determining locus (*r* < *R*).**

To clarify the parameter space under which neo-W alleles spread despite looser linkage with the selected locus (*R* > *r*), we focus on cases where dominance coefficients are equal in the two sexes, *h*^♀^ = *h*^♂^, and haploid selection only occurs in one sex (e.g., during male meiosis only). Table 3 then gives the conditions required for unlinked (*R* = 1/2) neo-W invasion when there is some ancestral sex-linkage (*r* < 1/2; e.g., the selected locus is on the ancestral sex chromosome and the novel sex-determining locus arises on an autosome). These special cases indicate that neo-W invasion occurs for a large fraction of the parameter space, even though the neo-W uncouples the sex-determining locus from a locus under selection. Fig 4 then demonstrates that under these conditions neo-W : alleles can spread when they are more loosely *or* more closely linked to the locus that experiences haploid selection, i.e., Conclusions 3B and 3C (c.f., Fig 3A for diploid sexually-antagonistic selection alone).

**Table 3.**
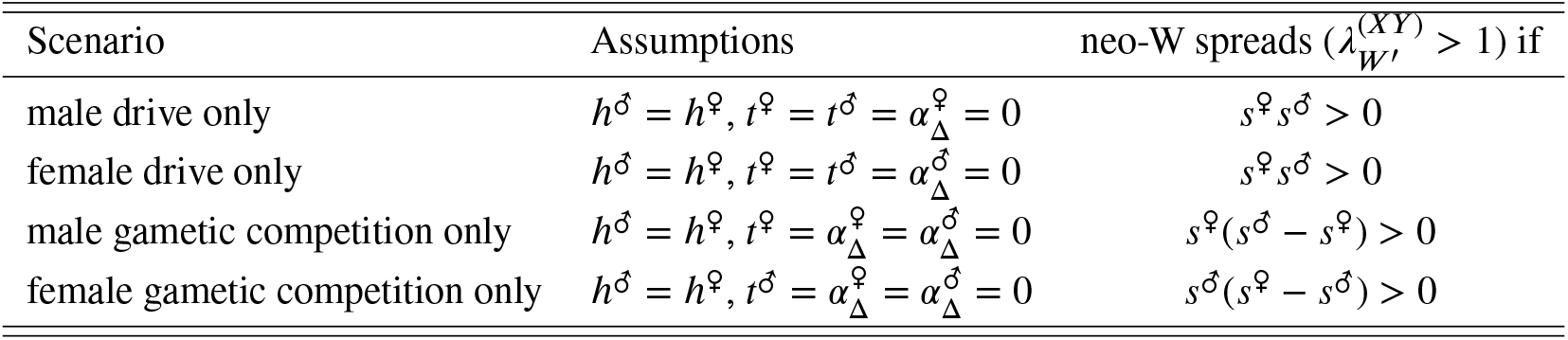
Invasion conditions for a neo-W allele at an unlinked locus (*R* = 1/2) into an ancestral XY system with linkage (*r* < 1/2) and a single form of haploid selection

We can also compare transitions in genetic sex-determination where sex-ratio bias increases, decreases, or remains equal. For example, if there is meiotic drive in males only 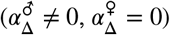, without gametic competition (*t*^♀^ = *t*^♂^ = 0) the zygotic sex ratio is initially biased only when the ancestral sex-determining system is XY (Figs 1B and 5A) and notZW (Figs 1B and 5B). If Fisherian sex-ratio selection were dominant, we would thus expect a difference in the potential for XY to ZW and ZW to XY transitions. However, invasion by a neo-W allele into an XY system and invasion by a neo-Y allele into a ZW system occur under the same conditions (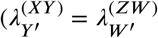 and 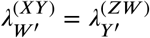, at least to order *ϵ*^2^), implying that,

*Conclusion 4:* **When selection is weak relative to recombination, trans-GSD transitions in the presence of haploid selection are favoured as often and as strongly whether they erase ancestral sex-ratio bias (benefiting from Fisherian sex ratio selection) or generate sex-ratio bias (benefiting from associations with selected alleles).**

For example, in Fig 5A neo-W alleles invade an ancestral-XY system where females are initially rare, equalizing the sex ratio (as occurs in [42]). However, Fig 5B shows that a neo-Y can invade the resulting ZW system under the same conditions. When *R* < 1/2, the invading neo-Y becomes associated with the male meiotic drive allele and the zygotic sex ratio evolves to become male-biased (as occurs in [43], beginning from ESD). In this case, the neo-Y spreads because it is often found in males and can, if it carries the driven allele *a*, benefit from haploid selection in males (Fig 5B).

**Fig 5.**
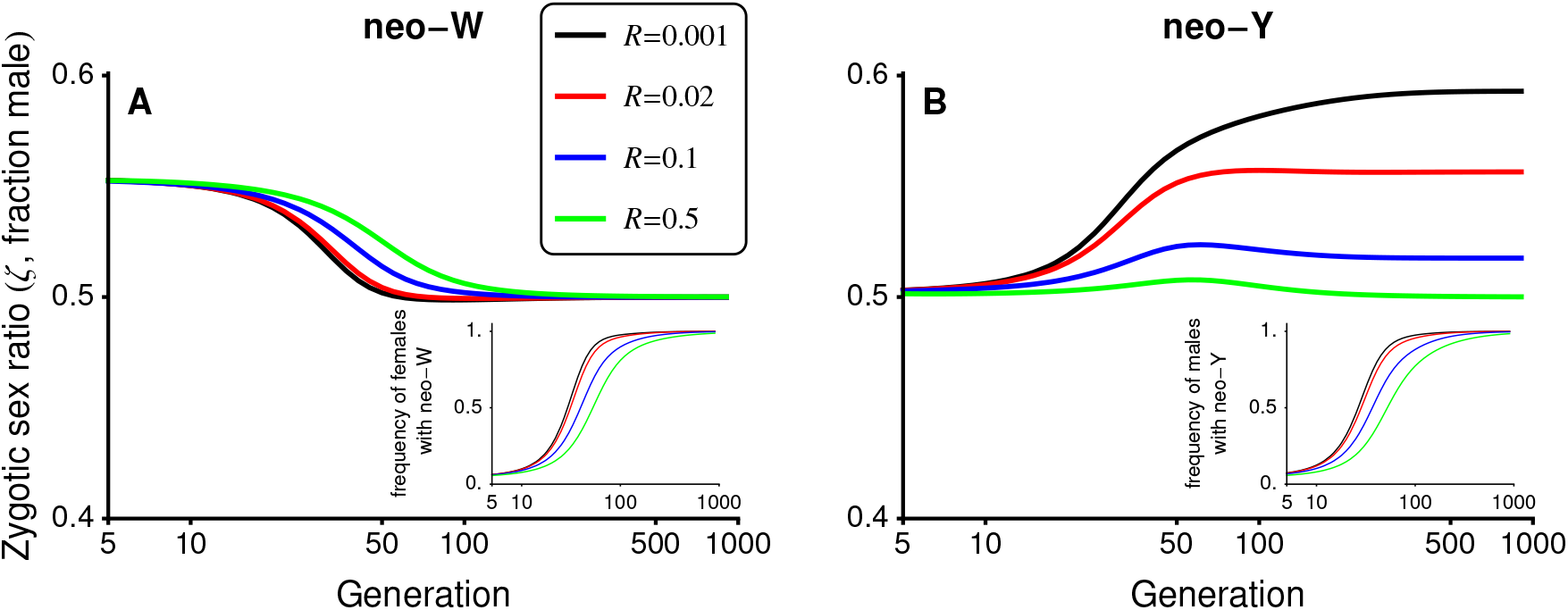
Fisherian sex-ratio selection alone is not a good predictor of turnover between sex-determining systems. In this figure, selection is ploidally antagonistic with haploid selection favouring the *a* allele during male meiosis. In panel A, male meiotic drive in an ancestral XY system causes a male bias (see Fig 1B), allowing a neo-W to invade and replace the ancestral sex-determining system (inset shows the frequency of females carrying a neo-W), which balances the zygotic sex ratio. In panel B, male drive in an ancestral ZW system has no effect on the zygotic sex ratio (50:50 at generation 0), yet a neo-Y can invade and replace the ancestral sex-determining system (inset shows the frequency of males carrying a neo-Y). Parameters: *s*^♀^ = *s*^♂^ = 0.2, *h*^♀^ = *h*^♂^ = 0.7, 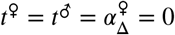, 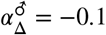, *r* = 0.02.

While equalizing the sex ratio and benefiting from associations with selected alleles are two primary reasons why haploid selection spurs sex chromosome transitions, more complex situations also arise. For example with *R* = 1/2 in Fig 5B (green curve), the neo-Y allele spreads despite the fact that it cannot benefit from drive because free recombination moves it randomly between driven and non-driven backgrounds. Nevertheless, the unlinked neo-Y can spread because males bearing it more often carry the non-driven allele *A* and have higher average diploid fitness compared to ZZ males, which bear a high frequency of the driven allele, *a*, from their mothers.

### Environmental sex determination

We next consider the case where the new sex-determining allele, *m*, causes sex to be determined probabilistically or by heterogeneous environmental conditions (environmental sex determination, ESD). In particular, we assume individuals carrying allele *m* develop as females with probability *k* ∈ (0, 1). In our deterministic model this means the fraction female in the subpopulation containing *m* is exactly *k*, even when *m* is rare (i.e., ESD does not introduce any additional variance in sex determination). We also assume that the environmental conditions that determine sex do not differentially affect the fitness of males versus females. Such correlations can favour environmental sex-determining systems by allowing each sex to be produced in the environment in which it has highest fitness; in the absence of these correlations previous theory would predict that ESD is favoured when it produces more equal sex ratios than the ancestral system (see reviews by [1, 31, 32]).

The characteristic polynomial determining the leading eigenvalue (equations S1.1) does not factor for ESD (0 < *k* < 1) as it does for a neo-Y (*k* = 0) or neo-W (*k* = 1) allele. We therefore focus on weak selection here, where the leading eigenvalue is

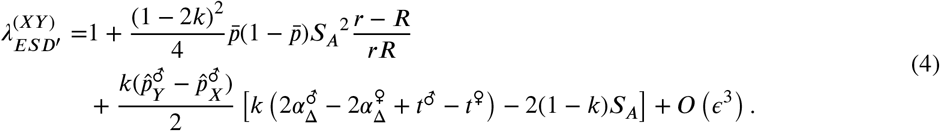

This reduces to 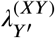 when *k* = 0 and 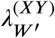 when *k* = 1.

Of particular interest are ESD mutations that cause half of their carriers to develop as females and half as males (*k* = 1/2), creating equal sex ratios. The spread of such mutations is determined by

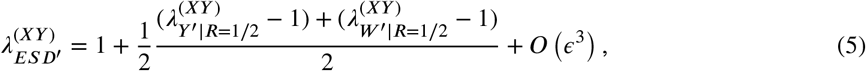

where 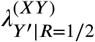 and 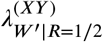 represent 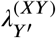 and 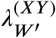 when evaluated at *R* = 1/2 (Equations 2 and 3). That is, ESD with *k* =1/2 behaves as if the **M** and **A** loci were unlinked, regardless of the actual value of *R*. This is because sex is randomized each generation in individuals bearing the *m* allele, preventing associations from building up between it and alleles at locus **A**. Eq (5) shows that the ESD mutation gets half of the fitness of a feminizing mutation (neo-W) and half of the fitness of a masculinizing mutation (neo-Y), but only has an effect one half of the time (the other half of the time it produces the same sex as the ancestral system would have). As discussed above, 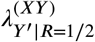 is necessarily less than or equal to one when selection is weak (Conclusion 3A), but 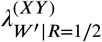 can be greater than one if there is haploid selection (see Conclusion 3C). That is, with haploid selection, an allele causing environmental-sex-determination can invade an ancestrally-XY system because it generates females that are either rare or have high fitness, in the same manner as a neo-W (likewise, ESD invades a ZW system for the same reasons that a neo-Y can).

Significantly, Eq (5) is the same whether ESD is invading an ancestrally XY or ZW system (because 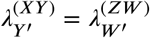 and 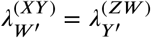). Thus, focusing solely on Fisherian selection to equalize the sex-ratio does not fully explain GSD to ESD transitions. For example, when the ancestral sex-determining system is XY the sex ratio is biased by male haploid selection. When the ancestral sex-determining system is ZW the sex ratio is not biased. Nevertheless, ESD is equally likely to invade both XY (through 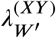) and ZW (through 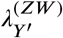) systems, equalizing the zygotic sex ratio in the former case but not in the latter. In addition, we note that ESD may not invade, even if the sex ratio is initially biased (e.g., with drive in males only, *r* < 1/2, *h*^♀^ = *h*^♂^, and *s*^♀^*s*^♂^ < 0, then 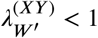, see Table 3). We conclude that, as with neo-W and neo-Y mutations:

*Conclusion 5:* **Transitions from genetic to environmental sex-determination are not straightforwardly predicted by selection to balance the zygotic sex ratio when haploid selection is present.**

## Discussion

New sex-determination systems are typically expected to spread when they equalise the sex ratio and/or when they increase linkage with loci that experience sex-differences in selection [33, 34]. In accordance with the latter mechanism, we find that sex-differences in selection at the haploid stage can favour cis- or trans-GSD transitions that tighten sex-linkage (Conclusion 3A &3B). Contrary to this expectation, however, we find that trans-GSD transitions can be favoured that loosen linkage with the sex-determining locus, either when linkage is initially tight (Conclusions 1 &#38; 2, Figs 2 & 3) or when there is haploid selection (Conclusion 3C, Figs 4 & 5). Furthermore, we show that the spread of new sex-determination systems is not dominated by selection to balance the sex ratio (Conclusions 4 & 5, Fig 5).

On the one hand, sex-ratio biases caused by haploid selection can facilitate trans-GSD transitions or transitions from genetic to environmental sex determination [42]. For instance, alleles favoured by haploid selection in males often become associated with the Y allele, which leads to an ancestral male-biased zygotic sex ratio. This male bias increases the potential for a neo-W or ESD allele to invade (Table 2), equalizing the sex ratio (e.g., see Fig 5B, for related examples see [42]). On the other hand, sex-ratio selection can be overwhelmed by additional selective effects, preventing a neo-W or ESD allele from invading, even if it would balance the sex ratio (e.g., when selection also acts in opposite directions in male and female diploids, Table 3). Indeed, transitions between sex-determining systems can generate stronger sex-ratio biases (e.g., Fig 5A and step 1 in [43]). Significantly, with weak selection, we find that there is no difference in conditions allowing XY to ZW and ZW to XY transitions (Conclusion 4) even when haploid selection always acts in the same sex (e.g., males). That is, the sex-ratio bias created by male haploid selection facilitates the spread of a neo-W allele into an XY system to the same degree that male haploid selection drives the spread of a neo-Y into a ZW system with a 1:1 sex ratio (Fig 5).

Because both Fisherian selection to equalize the sex ratio and the benefits of hitchhiking with driven alleles can facilitate transitions among sex chromosome systems, we predict that haploid selection should increase the lability of sex determination systems. Even in animal and plant species that have much larger and more conspicuous diploid phases than haploid phases, many loci have been shown to experience haploid selection through gamete competition and/or meiotic drive [38–41,51–56], which can generate biased sex-ratios [57–64]. In animals, a relatively small proportion of all genes are thought to be expressed and selected during competition in animal sperm [39, 65, 66]. Nevertheless, recent studies have demonstrated that sperm competition, even within a single ejaculate, can alter haploid allele frequencies and increase offspring fitness [67, 68]. Expression in the gamete is not required for haploid selection if the fitness of a gamete depends on its ability to condense DNA [69]. Furthermore, expression during gamete production often underlies systems of meiotic drive [70–72], which may be a common form of haploid selection in animals [73]. In plants, competition among gametophytes may be particularly important. It is estimated that 60-70% of all genes are expressed in the male gametophyte, and these genes exhibit stronger signatures of selection than randomly-chosen genes [74–76]. Furthermore, artificial selection pressures applied to male gametophytes are known to cause a response to selection (e.g., [77–80]).

Linking haploid expression with the evolution of sex-determination, a recent transcriptome analysis in *Rumex* shows that pollen-biased expression (relative to expression in flower buds or leaves) is enhanced among XY-linked genes compared to autosomal genes or compared to hemizygous genes that are only linked to the X [81]. In addition, Y-linked genes are over-expressed relative to X-linked genes in pollen (but not in flower buds or leaves). This suggests that the spread of neo-Y chromosomes in this clade could have been favoured through linkage with haploid selected genes rather than those under sexually antagonistic selection.

Frequent turnovers driven by haploid selection may help to explain the relative rarity of heteromorphic sex chromosomes in plants. If haploid selection is strong but selective differences between male and female diploids are weak, we specifically predict that trans-GSD transitions are favoured more strongly than cis-GSD transitions, with transitions to ESD intermediate (e.g., with 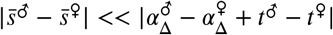 we have 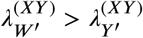; Eq 3). Among the relatively few dioecious clades in which multiple species have well characterized sex chromosomes [6], trans-GSD transitions have been inferred in *Silene* subsection *Otites* [15] and in *Salicaceae* [16, 17]. Assuming that transitions from dioecy to hermaphroditism (equal parental investment in male and female gametes) are favoured in a similar manner to the ESD examined here (equal probability of zygotes developing as males or females), our results suggest that competition among haploid pollen could drive transitions between dioecy and hermaphroditism, which are frequent in plants [82, 83]. To further examine this link, future theory could also include inbreeding, which is an important consideration during transitions between dioecy and hermaphroditism [84]. Future empirical studies could look for evidence of haploid selection acting on former sex chromosomes in hermaphroditic species (e.g., a study such as [81] on ancestral, rather than derived, sex chromosomes).

New sex-determining alleles have previously been shown to spread when they arise in linkage with loci that experience sex differences in selection because beneficial associations build up between alleles that determine sex and alleles that are favoured in that sex [35–37,43]. In support of this hypothesis, researchers have identified genes on recently derived sex chromosomes that might be under sexually-antagonistic selection [21,85–87]. However, we show that, if selected loci are tightly linked to the ancestral sex-determining locus, they can drive trans-GSD transitions that reduce sex-linkage (Conclusions 1 & 2), thus widening the range of genomic locations where selection could be driving observed trans-GSD transitions. In addition, we find that polymorphic sex determining systems (X, Y, and neo-W alleles all segregating) can be maintained when a selected locus is tightly linked to the ancestral sex-determining system (e.g., Fig S9B and Fig S9C), which is not possible with loose linkage [36]. This pair of conclusions apply in cases with or without haploid selection.

Our tight linkage result, in particular the prediction that invasion can lead to polymorphic sex determination, is consistent with empirical data from species in which new feminizing mutations are found segregating with ancestral XY loci. For example, in the platyfish (*Xiphophorus maculatus*), X,Y, and W alleles segregate at one locus (or two closely-linked loci) near to potentially sexually-antagonistic genes for pigmentation and sexual maturity [44,88–90]. Furthermore, several rodent species maintain feminizing alleles along with the ancestral X and Y sex-determination alleles (reviewed in [91]). In nine *Akadon* rodent species, it appears that male-determining-*sry* expression is suppressed by an autosomal feminizing allele (a neo-W allele), creating XY females [92, 93]. XY females have increased fitness relative to XX females [94]. However, it is not yet clear whether loci linked to the feminizing factor or the ancestral Y cause this effect. Most convincingly, in *Mus minutoides*, females can have XX, XX* or X*Y genotypes [95]. Previous theory would predict that the dominant X* chromosome (potentially an autosome that has fused with the sex chromosome) harbours female beneficial alleles, driving its spread. However, XX and XX* females have similar fitness, whereas X* Y female fitness is enhanced [96–98]. Although Y-linkage of female-beneficial alleles is counterintuitive, our model suggests that it can be stably maintained when linkage is initially tight between the sex determining region and the selected locus, subsequently favoring new feminizing mutations, which would be a parsimonious explanation for the spread of feminizing alleles in this case.

Our models assume that sex-determining alleles do not experience direct selection except via their associations with sex and selected alleles. However, in some cases, there may be significant degeneration around the sex-limited allele (Y or W) in the ancestral sex-determining region because recessive deleterious mutations and/or deletions accumulate in the surrounding non-recombining regions [99–102]. During trans-GSD transitions, but not cis-GSD transitions, any recessive deleterious alleles linked to the Y or W are revealed to selection in YY or WW individuals [4]. This phenomenon was studied by van Doorn and Kirkpatrick (2010) [36], who found that degeneration can prevent fixation of a neo-W or a neo-Y allele, leading to a mixed sex-determining system where the ancestral and new sex-determining loci are both segregating. However, they noted that very rare recombination events around the ancestral sex-determining locus can allow the completion of trans-GSD transitions. Degeneration around the Y or W could explain why trans-GSD transitions are not observed to be much more common than cis-GSD transitions despite the fact that our models demonstrate that they are favoured under a wider range of conditions, especially with haploid selection. For example, there are a dozen sex chromosome configurations among Dipteran species but only one transition between male and female heterogamety [9], but Y degeneration or absence is also very common among *Diptera* [9].

In this study, we have only considered new sex-determining alleles of large effect. However, we expect similar selective forces to act on masculinizing/feminizing alleles of weaker effect. For example, small effect masculinizing/feminizing alleles within a threshold model of sex determination can be favoured when linked to loci that experience sexually-antagonistic selection [37]. These results echo those for large-effect neo-Y/neo-W alleles [35, 36]. It should be noted, however, that the dynamics of sex-determining alleles with very weak effect will be influenced by genetic drift, which itself has been shown to bias transitions towards epistatically-dominant sex-determining systems when there is no direct selection [103].

## Conclusion

We have shown that tight sex-linkage and haploid selection can drive previously unexpected transitions between sex-determining systems. In particular, both can select for new sex-determining loci that are more loosely linked to loci under selection (Conclusions 2 & 3C). In addition, haploid selection can cause transitions in GSD analogous to those caused by purely sexually-antagonistic selection, eliminating the need for differences in selection between male and female diploids (Conclusion 3A, 3B & 3C). We conclude that haploid selection should be considered as a pivotal factor driving transitions between sex-determining systems. Further, transitions involving haploid selection can eliminate or generate sex-ratio biases; to leading order, selection to balance the sex ratio and the benefits of hitch-hiking with haploid selected alleles, leading to a biased sex ratio, are of equal magnitude (Conclusions 4 & 5). Overall, our results suggest several novel scenarios under which new sex-determining systems are favoured, which could help to explain why the evolution of sex-determining systems is so dynamic.

## Acknowledgments

We thank Georgy Sandler and Stephen Wright for sharing their results with us, and we thank Bret Payseur and three anonymous reviewers for helpful comments on this manuscript.

## Supporting information

### Supplementary Files

**S1 File. Supplementary *Mathematica* file.** This file can be used to re-derive our results and generate figures. Please contact authors for access (or for a PDF version).

## S1 Appendix Recursion equations

In each generation we census the genotype frequencies in male and female gametes/gametophytes (hereafter, gametes) between meiosis (and any meiotic drive) and gametic competition. At this stage we denote the frequencies of X- and Y-bearing gametes from males and females 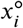 and 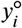. The superscript ∘ ∈ {♂, ♀} specifies the sex of the diploid that the gamete came from. The subscript *i* ∈ {1,2,3,4} specifies the genotype at the selected locus **A** and at the novel sex-determining locus **M**, where 1 = *AM*, 2 = *aM*, 3 = *Am*, and 4 = *am*. The gamete frequencies from each sex sum to one, 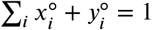.

Competition then occurs among gametes of the same sex (e.g., among eggs and among sperm separately) according to the genotype at the **A** locus (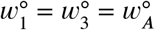, 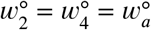, see Table 1). The genotype frequencies after gametic competition are 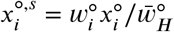 and 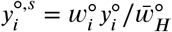, where 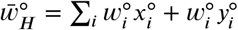 is the mean fitness of male (∘ = ♂) or female (∘ = ♀) gametes.

Random mating then occurs between gametes to produce diploid zygotes. The frequencies of XX zygotes are then denoted as *xx_ij_*, XY zygotes as *xy_ij_*, and YY zygotes as *yy_ij_*, where **A** and **M** locus genotypes are given by *i,j* ∈ {1,2,3,4}, as above. In XY zygotes, the haplotype inherited from an X-bearing gamete is given by *i* and the haplotype from a Y-bearing gamete is given by *j*. In XX and YY zygotes, individuals with diploid genotype *ij* are equivalent to those with diploid genotype *ji*; for simplicity, we use *xx_ij_* and *yy_ij_* with *i* ≠ *j* to denote the average of these frequencies, 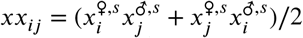 and 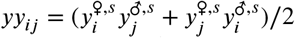.

Denoting the **M** locus genotype by *b* ∈ { ***MM***, ***M**m*, *mm* } and the **X** locus genotype by *c* ∈ { *XX, XY, YY* }, zygotes develop as females with probability *k_bc_*. Therefore, the frequencies of XX females are given by 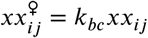, XY females are given by 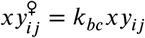, and YY females are given by 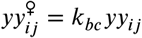. Similarly, XX male frequencies are 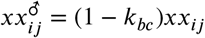, XY male frequencies are 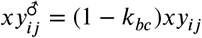, and YY males frequencies are 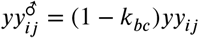. This notation allows both the ancestral and novel sex-determining regions to determine zygotic sex according to an XY system, a ZW system, or an environmental sex-determining system. In addition, we can consider any epistatic dominance relationship between the two sex-determining loci. Here, we assume that the ancestral sex-determining system (**X** locus) is XY (*k_MMXX_* = 1 and *k_MMXY_* = *k_MMYY_* = 0) or ZW (*k_MMZZ_* = 0 and *k_MMZW_* = *k_MMWW_* = 1) and epistatically recessive to a dominant novel sex-determining **M** (*k_Mmc_* = *k_mmc_* = *k*).

Selection among diploids then occurs according to the diploid genotype at the **A** locus, *l* ∈ { *AA*, *Aa*, *aa* }, for an individual of type *ij* (see Table 1). The diploid frequencies after selection in sex ∘ are given by 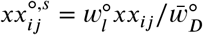, 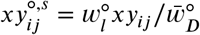, and 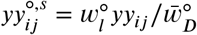, where 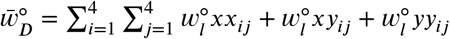 is the mean fitness of diploids of sex ∘.

Finally, these diploids undergo meiosis to produce the next generation of gametes. Recombination and sex-specific meiotic drive occur during meiosis. Here, we allow any relative locations for the **X**, **A**, and **M** loci by using three parameters to describe the recombination rates between them. *R* is the recombination rate between the **A** and **M** loci, *ρ* is the recombination rate between the **M** and **X** loci, and *r* is the recombination rate between the **A** and **X** loci (Fig 1). Table S1 shows replacements that can be made for each possible ordering of the loci assuming that there is no cross-over interference. During meiosis in sex ∘, meiotic drive occurs such that, in *Aa* heterozygotes, a fraction *α*^∘^ of gametes produced carry the *A* allele and (1 - *α*^∘^) carry the *a* allele.

Among gametes from sex ∘, the frequencies of haplotypes (before gametic competition) in the next generation are given by

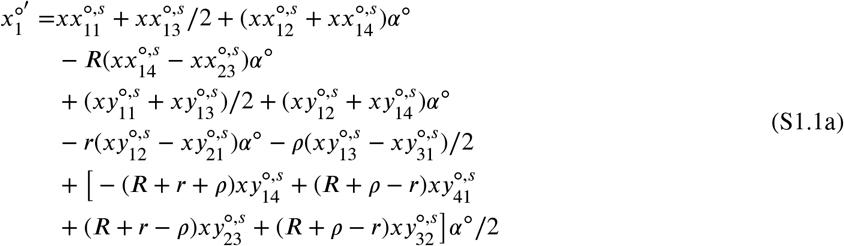

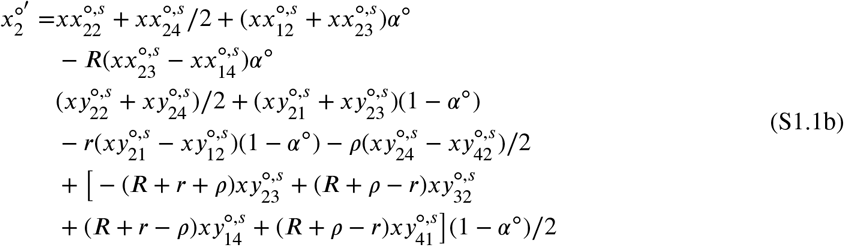

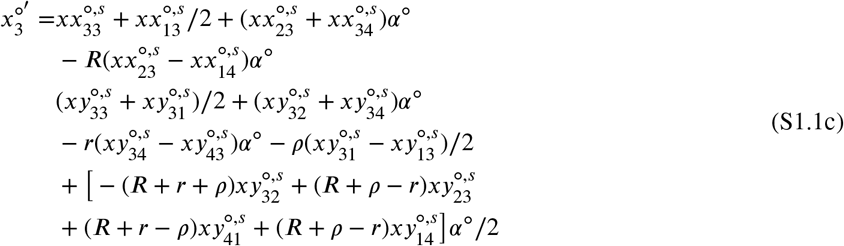

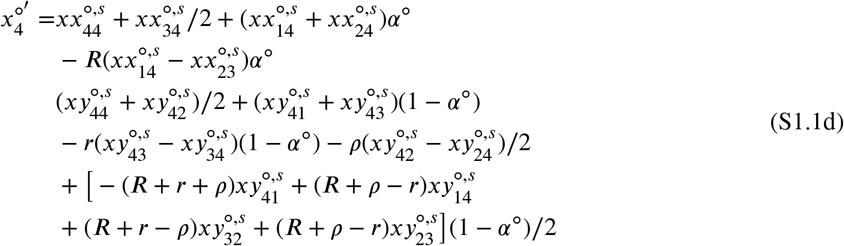

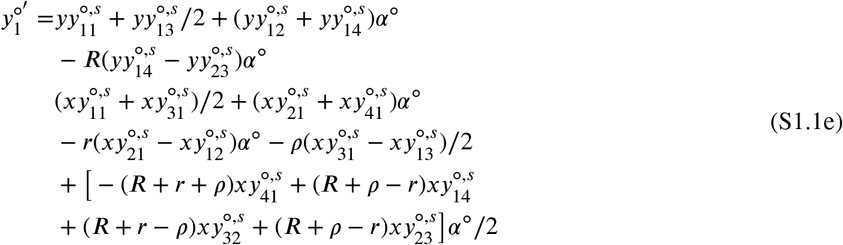

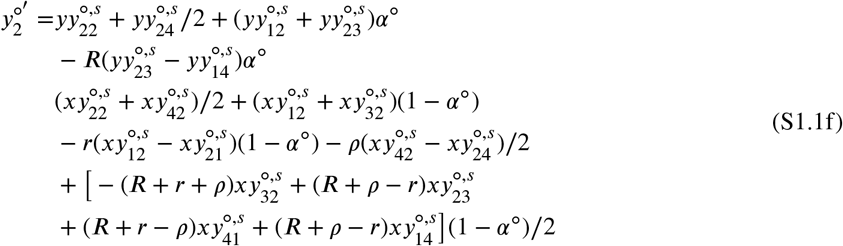

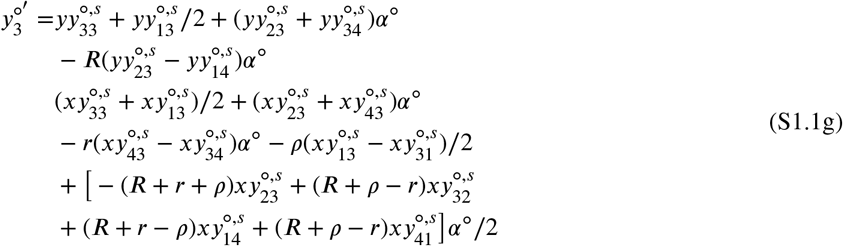

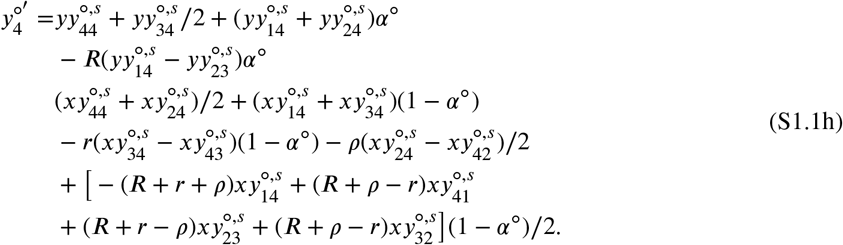

The full system is therefore described by 16 recurrence equations (three diallelic loci in two sexes, 2^3^ × 2 = 16). However, not all diploid types are produced under certain sex-determining systems. For example, with the *M* allele fixed and an ancestral XY sex-determining system, there are XX females and XY males 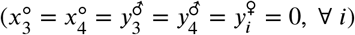. In this case, the system only involves six recursion equations, which we assume below to calculate the equilibria.

## S2 Appendix Resident equilibria and stability

In the resident population (allele ***M*** fixed), we follow the frequency of *A* in X-bearing female gametes, 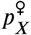, and X-bearing male gametes, 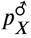, and Y-bearing male gametes, 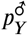. We also track the total frequency of Y among male gametes, *q*, which may deviate from 1/2 due to meiotic drive in males. These four variables determine the frequencies of the six resident gamete types, which sum to one in each sex: 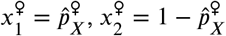, 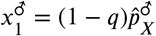, 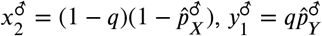, and 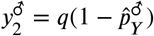. Mean fitnesses in the resident population are given in Table S2.

Various forms of selection can maintain a polymorphism at the **A** locus, including sexually antagonistic selection, overdominance, conflicts between diploid selection and selection upon haploid genotypes (ploidally-antagonsitic selection, [48]), or a combination of these selective regimes (see below).

In particular special cases, e.g., no sex-differences in selection or meiotic drive (*s*^♂^ = *s*^♀^, *h*^♂^ = *h*^♀^, and *α*^♂^ = *α*^♀^ = 1/2), the equilibrium allele frequency and stability can be calculated analytically without assuming anything about the relative strengths of selection and recombination. However, here, we focus on two regimes (tight linkage between **A** and **X** and weak selection) in order to make fewer assumptions about fitnesses.

### Tight linkage between X and A loci

We first calculate the equilibria in the ancestral population when the recombination rate between the **X** and **A** loci is small (*r* of order *ϵ*). Selection at the **A** locus will not affect evolution at the novel sex-determining locus, **M**, if one allele is fixed on all backgrounds. We therefore focus on the five equilibria that maintain both *A* and *a* alleles, four of which are given to leading order by:

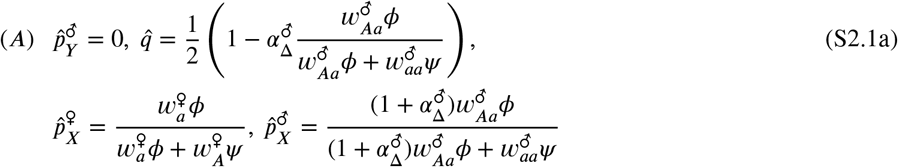

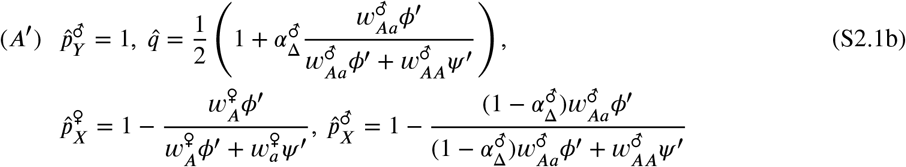

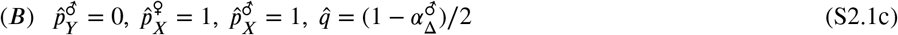

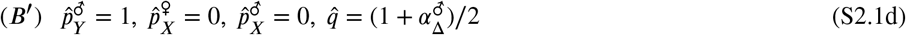

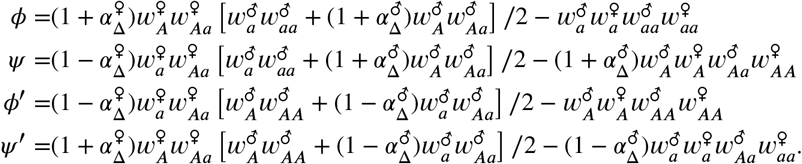

A fifth equilibrium (*C*) also exists where *A* is present at an intermediate frequency on the Y background 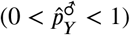. However, equilibrium (*C*) is never locally stable when *r* ≈ 0 and is therefore not considered further. Thus, the Y background can either be fixed for the *a* allele (equilibria (*A*) and (*B*)) or the *A* allele (equilibria (*A′*) and (*B′*)). The X background can then either be polymorphic (equilibria (*A*) and (*A′*)) or fixed for the alternative allele (equilibria (*B*) and (*B′*)). Since equilibria (*A*) and (*B*) are equivalent to equilibria (*A′*) and (*B′*) with the labelling of *A* and *a* alleles interchanged, we discuss only equilibria (*A*) and (*B*), in which the Y background is fixed for the *a* allele. If there is no haploid selection 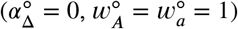, these equilibria are equivalent to those found by [49] and [50].

We next calculate when equilibria (*A*) and (*B*) are locally stable for *r* = 0. According to the ‘small parameter theory’ [104, 105], these stability properties are unaffected by small amounts of recombination between the sex-determining locus and the **A** locus, although equilibrium frequencies may be slightly altered. For the *a* allele to be stably fixed on the Y background we need 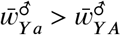 where 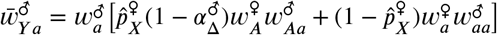 and 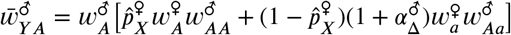. That is, Y-*a* haplotypes must have higher fitness than Y-*A* haplotypes. Substituting in 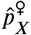 from Eq (S2.1), fixation of the *a* allele on the Y background requires that *γ_i_* > 0 where 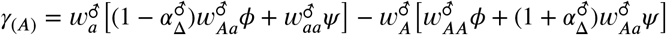 for equilibrium (*A*) and 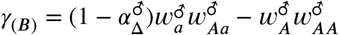 for equilibrium (*B*). Stability of a polymorphism on the X background (equilibrium (*A*)) further requires that *ϕ* > 0 and *ψ* > 0. Fixation of the *a* allele on the X background (equilibrium (*B*)) can be stable only if equilibrium (*A*) is not, as it requires *ψ* < 0.

### Selection weak relative to recombination

Here, we assume that selection is weak relative to recombination (*s*^∘^, *t*^∘^, 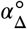 of order *ϵ*). The maintenance of a polymorphism at the **A** locus then requires that

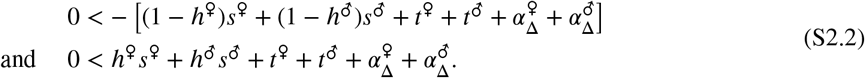

which indicates that a polymorphism can be maintained by various selective regimes.

Given that a polymorphism is maintained at the **A** locus by weak selection, the frequencies of *A* in each type of gamete are the same 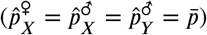 and given, to leading order, by

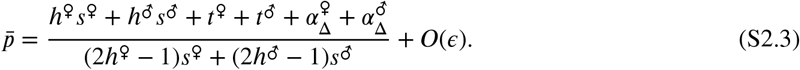

Differences in frequency between gamete types are of *O*(*ϵ*):

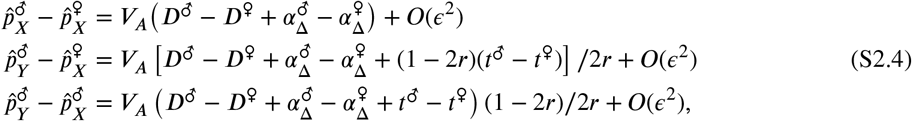

where 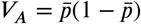 is the variance in the frequency of *A* and 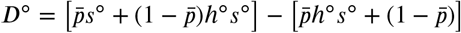 corresponds to the difference in fitness between *A* and *a* alleles in diploids of sex ∘ ∈ { ♀, ♂ }. The frequency of Y-bearing male gametes depends upon the difference in the frequency of the *A* allele between X- and Y-bearing male gametes and the strength of meiotic drive in favour of the *A* allele in males, 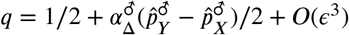. Without gametic competition or drive 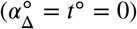 our results reduce to those of [35].

## S3 Appendix Invasion conditions

A rare sex-determining allele, *m*, will spread in an XY sex-determining system when the leading eigenvalue, 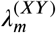, of the Jacobian matrix derived from the eight mutant recursion equations (given by S1.1c,d,g,h), evaluated at the ancestral equilibrium, is greater than one. Because a neo-Y (neo-W) is always in males (females) and is epistatically dominant to the ancestral sex-determining locus, the characteristic polynomial factors into two quadratics times 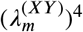, which greatly simplifies analysis. One quadratic governs the change in the frequency of the new sex determination factor associated with the *A* and *a* allele frequencies, summing across backgrounds at the old sex-determining locus (i.e., summing haplotypes bearing the original X and Y alleles). The second quadratic describes the dynamics of the difference in the *m-A* and *m-a* haplotypes between X and Y backgrounds. Asymptotically, this difference is constrained to grow at a rate equal to or less than the sum while the *m* allele is rare, because the frequencies can never become negative. We therefore focus on the leading eigenvalue, 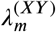, which is the largest root of the quadratic *f*(*x*) = *x*^2^ + *bx* + *c* = 0, where 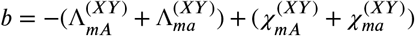 and 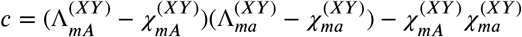 (details in S1 File), see Table 2.

When *R* = 0 the two roots are 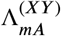 and 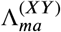, and the leading eigenvalue is the larger of the two. When *R* > 0 then 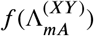 and 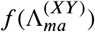 are of opposite signs, and the leading eigenvalue must fall between these two quantities (details in S1 File). Thus, 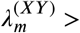 if both 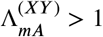 and 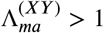; similarly, 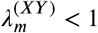 if both 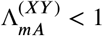 and 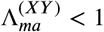. If only one haplotypic growth rate is greater than one (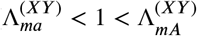 or 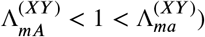), then 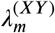 is greater than one when condition (1) is met. Thus, the invasion of a new sex-determining allele is determined by the haplotypic growth rates (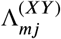 terms), which do not account for loss due to recombination, and the dissociative force that breaks apart these haplotypes by recombination 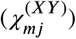. For tight linkage between the ancestral sex-determining locus (**X**) and the selected locus (**A**) we can calculate these terms explicitly (see below). For weak selection we approximate the leading eigenvalue with a Taylor series. The leading eigenvalue for any *k* is given up to order *ϵ^2^* by Eq (4).

### Tight linkage between A and X (*r* ≈ 0)

Here, we explore the conditions under which a neo-W invades an XY system assuming that the **A** locus is initially in tight linkage with the ancestral sex-determining locus (*r* ≈ 0). We disregard neo-Y mutations, which never spread given that the ancestral population is at a stable equilibrium (see S1 File for proof).

Starting with the simpler (*B*) equilibrium, the haplotypic growth rates 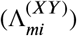 and dissociative forces 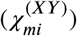 are

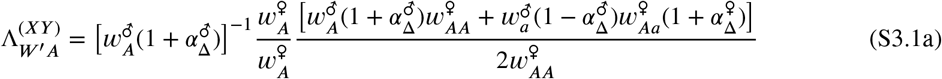

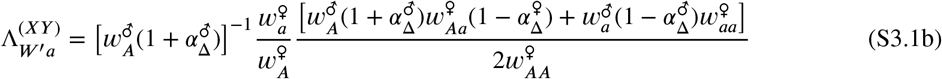

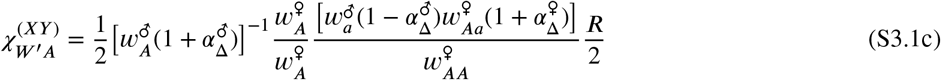

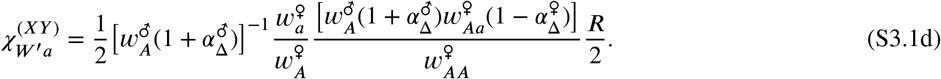

Haploid selection impacts the spread of neo-W haplotypes in three ways (also seen in Table 2). Firstly, the zygotic sex ratio becomes male biased, *ζ* > 1/2, when the *a* allele (which is fixed on the Y) is favoured during competition among male gametes or by meiotic drive in males. Specifically, at equilibrium (*B*), female zygote frequency is 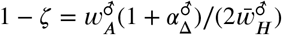 where 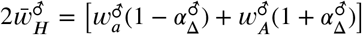 has been canceled out in equations (S3.1) to leave the term 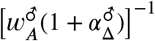. Male biased sex ratios facilitate the spread of a neo-W because neo-W alleles cause the zygotes that carry them to develop as the rarer, female, sex.

Secondly, haploid selection in females selects on neo-W haplotypes directly. At equilibrium (*B*), the fitness of female gametes under the ancestral sex-determining system is 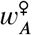 such that the relative fitnesses of neo-W-*A* and neo-W-*a* haplotypes during female gametic competition are 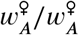 and 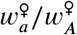 (see terms in equation S3.1). Meiotic drive in females will also change the proportion of gametes that carry the *A* versus *a* alleles, which will be produced by heterozygous females in proportions 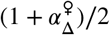 and 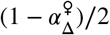, respectively. These terms are only associated with heterozygous females, i.e., they are found alongside 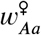.

Thirdly, haploid selection in males affects the diploid genotypes of females by altering the allele frequencies in the male gametes with which female gametes pair. At equlibrium (*B*), neo-W female gametes will mate with X-*A* male gametes with probability 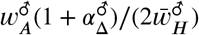 and Y-*a* male gametes with probability 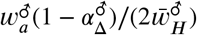, where the 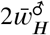 terms have been canceled in Eq (S3.1) (as mentioned above). Thus, neo-W-*A* haplotypes are found in *AA* female diploids with probability 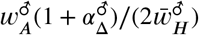 (e.g., first term in square brackets in the numerator of equation S3.1a) and in *Aa* female diploids with probability 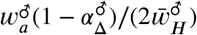 (e.g., the second term in square brackets in the numerator of equation S3.1a).

The other terms in equations (S3.1) are more easily interpreted if we assume that there is no haploid selection in either sex, in which case 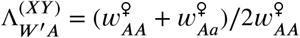 and 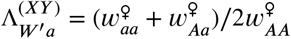. Neither haplotype can spread under purely sexually-antagonistic selection at equilibrium (*B*), with directional selection in each sex. Essentially, the X is then already as specialized as possible for the female beneficial allele (*A* is fixed on the X background), and the neo-W often makes daughters with the Y-a haplotype, increasing the flow of *a* alleles into females, which reduces the fitness of those females.

If selection doesn’t uniformly favour *A* in females, however, neo-W-*A* haplotypes and/or neo-W-*a* haplotypes can spread (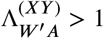 and/or 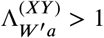). A neo-W-A haplotype can spread 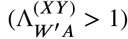 when 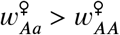, despite the fact that a neo-W brings Y-*a* haplotypes into females. In this case the *a* aAlele is favoured by selection in females despite *A* being fixed on the X background. For this equilibrium to be stable (i.e., to keep *A* fixed on the X), X-*a* cannot be overly favoured in females and X-*A* must be sufficiently favoured in males (for example, by overdominance in males). Specifically, from the stability conditions for equilibrium (*B*), we must have 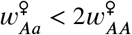 and 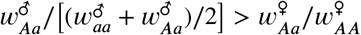.

Still considering 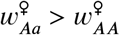, the neo-W can also spread alongside the *a* allele 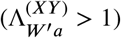 if 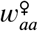 is large enough such that 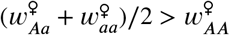. This can occur with overdominance or directional selection for *a* in females (Fig 2B,C). In this case, *a* is favoured on the ancestral Y background in males and on the ancestral X background in females (comparing *Aa* to *AA* genotypes in females) but *A* is fixed on the X background due to selection in males. The neo-W-a haplotype can spread because it produces females with higher fitness *Aa* and *aa* genotypes.

Similar equations can be derived for equilibrium (A) by substituting the equilibrium frequencies into Table 2

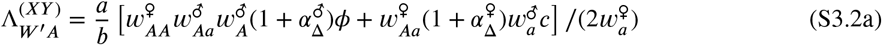

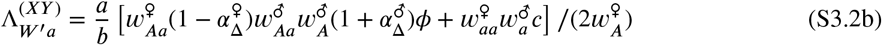

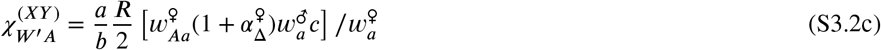

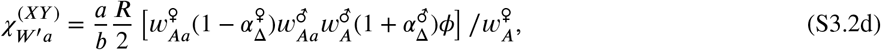

where

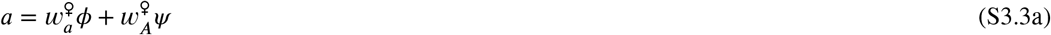

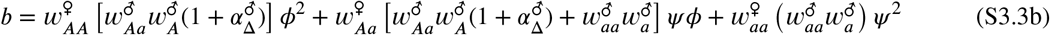

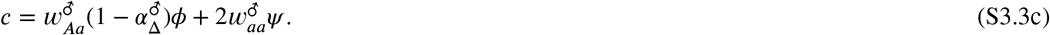

As with equilibrium (**B**), haploid selection again modifies invasion fitnesses by altering the ancestral sex ratio, *ζ* (see Table 2), and directly selecting upon female gametes, through 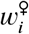. The only difference is that resident XX females are no longer always homozygote *AA* and males are no longer always heterozygote *Aa*. Thus the effect of haploid selection in males is reduced, as is the difference in fitness between neo-W haplotypes and resident X haplotypes, as both can be on any diploid or haploid background.

The other terms are easier to interpret in the absence of haploid selection. For instance, without haploid selection, the neo-W-*A* haplotype spreads 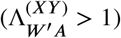 if and only if

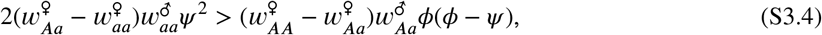

where 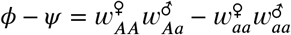 and both *φ* and *ψ* are positive when equilibrium (A) is stable. In contrast to equilibrium (B), a neo-W haplotype can spread under purely sexually-antagonistic selection (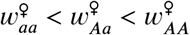 and 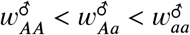). The neo-W-A can spread as long as it becomes associated with females that bear more *A* alleles than observed at equilibrium (A), effectively specializing on female fitness.

Without haploid selection, the neo-W-*a* haplotype spreads 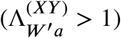 if and only if

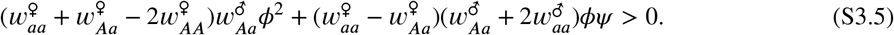

This condition cannot be met with purely sexually antagonistic selection (as both terms on the left-hand side would then be negative), but it can be met under other circumstances. For example, with overdominance in males there is selection for increased *A* frequencies on the X background in males, which are always paired with Y-*a* haplotypes. Directional selection for *a* in females can then maintain a polymorphism at the **A** locus on the X background. This scenario selects for a modifier that increases recombination between the sex chromosomes (e.g., blue region of Fig 2d in [50]) and facilitates the spread of neo-W-*a* haplotypes, which create females bearing more *a* alleles than the ancestral X haplotype does.

In absence of haploid selection, the fact that a less closely linked neo-W (R > 0) can invade an XY system with tight sex-linkage can also be reached from Equation 7 in [36]; for example, with no polymorphism on the Y (*V_Y_* = 0) and an allelic substitution favoured in females (*α^f^*, 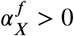) a loosely linked neo-W can invade given the allelic substitution is sufficiently disfavoured on the X in males 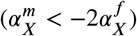, although it is unclear from their implicit equation if and when such an equilibrium is stable.

### Role of haploid selection with tight linkage between X and A loci

Haploid selection generally expands the conditions under which neo-W alleles can spread within ancestral systems that have evolved tight linkage between the sex-determining locus and a selected locus (*r* ≈ 0). First, haploid selection can allow a polymorphism to be maintained when it would not under diploid selection alone (e.g., with directional selection in diploids). In cases of ploidally-antagonistic selection, where there is a balance between alleles favored in the haploid stage and the diploid stage, neo-W alleles - even if unlinked to the selected locus - can spread (Fig S8). Second, even when diploid selection could itself maintain a polymorphism, haploid selection can increase the conditions under which transitions among sex-determining systems are possible. Of particularly importance, when selection is sexually-antagonistic in diploids (*s*^♀^*s*^♂^ < 0 and 0 < *h*^∘^ < 1), an unlinked neo-W (*R* =1/2) cannot invade unless there is also haploid selection (see proof in S1 File; Fig 3 and Fig S3). More generally, haploid selection alters the conditions under which neo-W alleles can spread (compare Fig S4-Fig S7 to Fig 2).

Male haploid selection in favour of the *a* allele 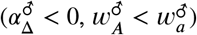 generates male-biased sex ratios at equilibria (A) and (B), where Y-*a* is fixed 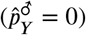. Male-biased sex ratios facilitate the spread of neo-W-*A* and neo-W-*a* haplotypes (increasing 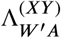 and 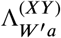). Panels A-C in Fig S4 and S5 show that neo-W haplotypes tend to spread for a wider range of parameters when sex ratios are male biased, compared to Fig 2 without haploid selection. By contrast, male haploid selection in favour of the *A* allele generates female-biased sex ratios and reduces 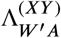 and 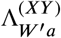), as demonstrated by panels D-F in Fig S4 and Fig S5.

Female heploid selection generates direct selection on the neo-W-*A* and neo-W-*a* haplotypes as they spread in females. Thus, female haploid selection in favour of the *a* allele tends to increase 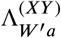 and decrease 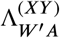, as shown by panels A-C in Fig S6 and Fig S7. Conversely, female haploid selection in favour of the *A* allele increases 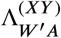 and decreases 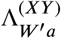), see panels D-F in Fig S6 and Fig S7.

Thus, the impact of haploid selection on transitions between sex-determining systems must be considered as two sides of a coin: it can generate sex ratio biases that promote transitions that equalize the sex ratio, but it can also direct select for transitions that cause sex ratios to become biased.

## Supplementary Tables

**Table S1.** Substitutions for different loci orders assuming no interference.

| Order of loci |  |
| --- | --- |
| SDR-A-M | $\rho = r(1 - R) + R(1 - r)$ |
| SDR-M-A | $r = \rho(1 - R) + R(1 - \rho)$ |
| A-SDR-M | $R = r(1 - \rho) + \rho(1 - r)$ |

**Table S2.** Mean fitnesses and zygotic sex ratio in the resident population (*M* fixed, XY sex determination).

| Sex & Life Cycle Stage | Mean Fitness |
| --- | --- |
| female gametes ($\bar{w}_H^\varnothing$) | $p_X^\varnothing w_A^\varnothing + (1 - p_X^\varnothing) w_a^\varnothing$ |
| X-bearing male gametes ($\bar{w}_{HX}^\delta$) | $p_X^\delta w_A^\delta + (1 - p_X^\delta) w_a^\delta$ |
| Y-bearing male gametes ($\bar{w}_{HY}^\delta$) | $p_Y^\delta w_A^\delta + (1 - p_Y^\delta) w_a^\delta$ |
| male gametes ($\bar{w}_H^\delta$) | $(1 - q) \bar{w}_{HX}^\delta + q \bar{w}_{HY}^\delta$ |
| females ($\bar{w}^\varnothing$) | $\frac{[p_X^\varnothing w_A^\varnothing p_X^\delta w_A^\delta w_{AA}^\varnothing + (1 - p_X^\varnothing) w_a^\varnothing p_X^\delta w_A^\delta w_{Aa}^\varnothing + p_X^\varnothing w_A^\varnothing (1 - p_X^\delta) w_a^\delta w_{Aa}^\varnothing + (1 - p_X^\varnothing) w_a^\varnothing (1 - p_X^\delta) w_a^\delta w_{aa}^\varnothing]}{(\bar{w}_H^\varnothing \bar{w}_{HX}^\delta)}$ |
| males ($\bar{w}^\delta$) | $\frac{[p_X^\varnothing w_A^\varnothing p_Y^\delta w_A^\delta w_{AA}^\delta + (1 - p_X^\varnothing) w_a^\varnothing p_Y^\delta w_A^\delta w_{Aa}^\delta + p_X^\varnothing w_A^\varnothing (1 - p_Y^\delta) w_a^\delta w_{Aa}^\delta + (1 - p_X^\varnothing) w_a^\varnothing (1 - p_Y^\delta) w_a^\delta w_{aa}^\delta]}{(\bar{w}_H^\varnothing \bar{w}_{HY}^\delta)}$ |
| fraction zygotes male ($\zeta$) | $q [p_Y^\delta w_A^\delta + (1 - p_Y^\delta) w_a^\delta] / \bar{w}_H^\delta$ |

## Supplementary Figures

**Fig S1.**
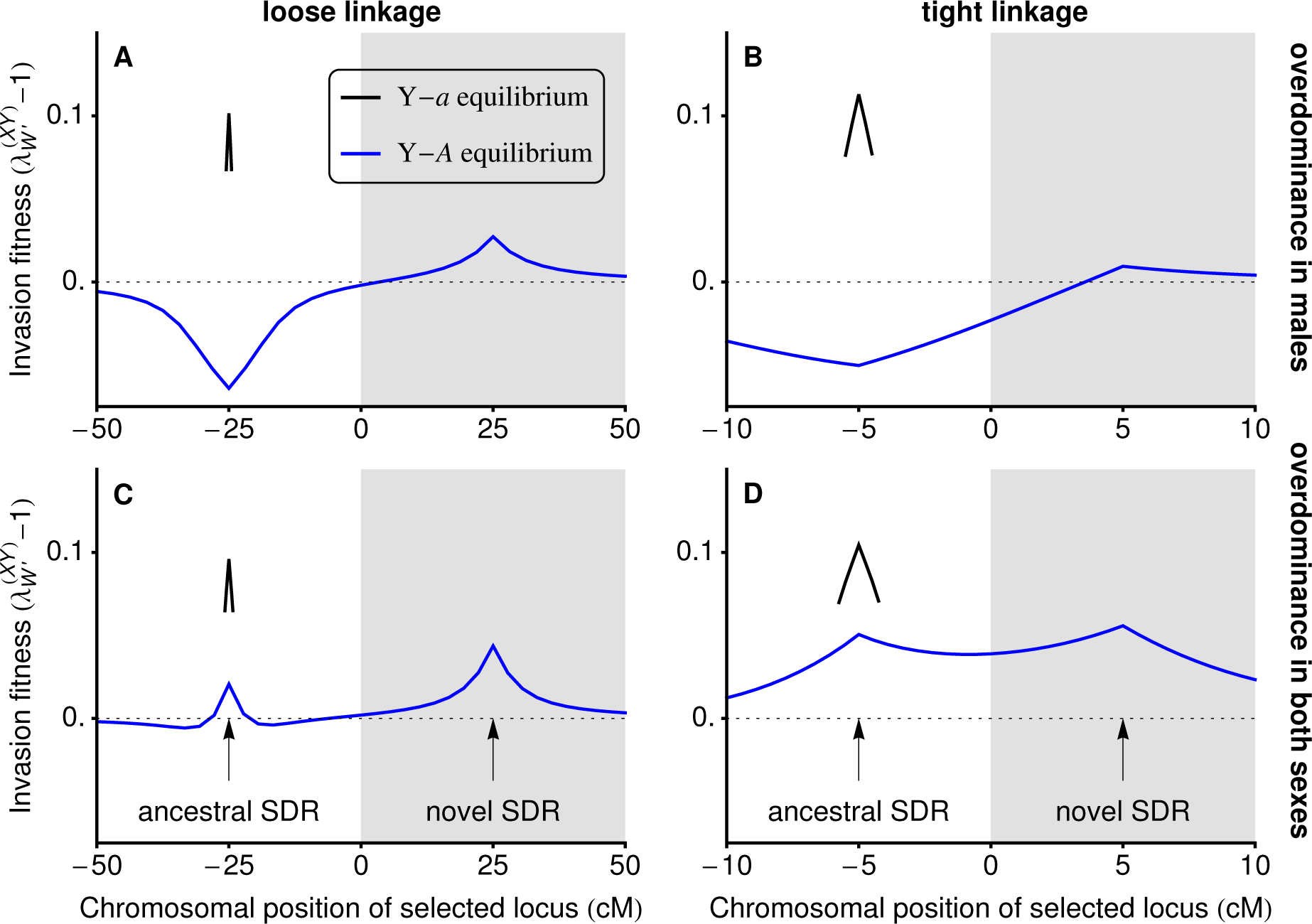
With overdominance, loci near to the ancestral sex-determining locus (*r* ≈ 0) can favour neo-W alleles that are less tightly linked (*R* > *r*). In panels A and B, the *a* allele is favoured in females 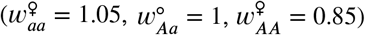 and selection in males is overdominant 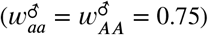. In panels C and D, selection in males and females is overdominant 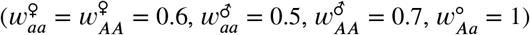. There is no haploid selection 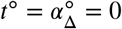. These parameters are marked by daggers in Fig 2B and C, which show that neo-W invasion is expected for any 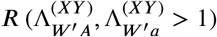 if the *a* allele is nearly fixed on the Y (black lines in this figure; not stable for *r* ≫ 0). Equilibria where the *A* allele is more common among Y-bearing male gametes can also be stable and allow neo-W invasion for these parameters (blue lines).

**Fig S2.**
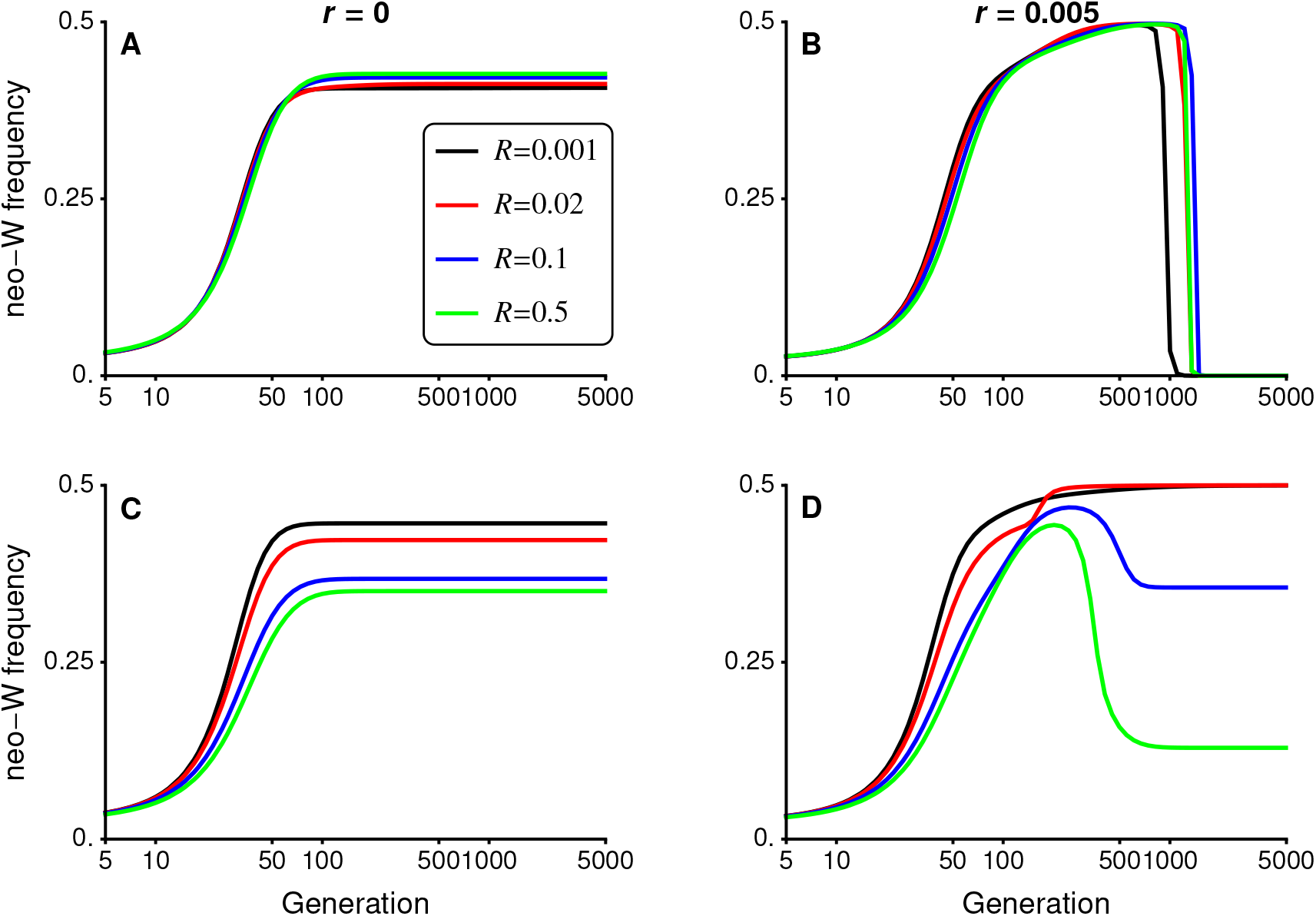
Following invasion by a neo-W allele, there can be a complete transition to a new sex-determining system, maintenance of both ancestral-XY and neo-ZW sex determining systems, or loss of the new sex-determining allele. Here, we plot the frequency of the neo-W allele among female gametes. Panels A, C and D show cases where a steady state is reached with the neo-W at a frequency below 0.5, in which case ancestral-X and Y alleles also both segregate. In all cases, we assume that the *a* allele is initially more common than the *A* allele on the Y background (Y-a is fixed when *r* = 0). When *r* > 0 (panels B and D), Y-*A* haplotypes created by recombination can become more common than Y-a haplotypes as the neo-W spreads. In B, this leads to loss of the neo-W and the system goes to an equilibrium with X-*a* and Y-*A* haplotypes fixed (equilibrium *A′*), such that all females have the high fitness genotype *aa* and all males are *Aa*. For the parameters in B, neo-W alleles have negative invasion fitness when the Y-*A* haplotype is ancestrally more common than Y-*a* (compare blue to black curves in Fig S1A and Fig S1B near the ancestral sex-determining locus). In contrast, the neo-W is not lost in panel D as it is favoured regardless of whether Y-*A* or Y-*a* haplotypes predominate (again, compare blue to black curves in Fig S1C and Fig S1D).

**Fig S3.**
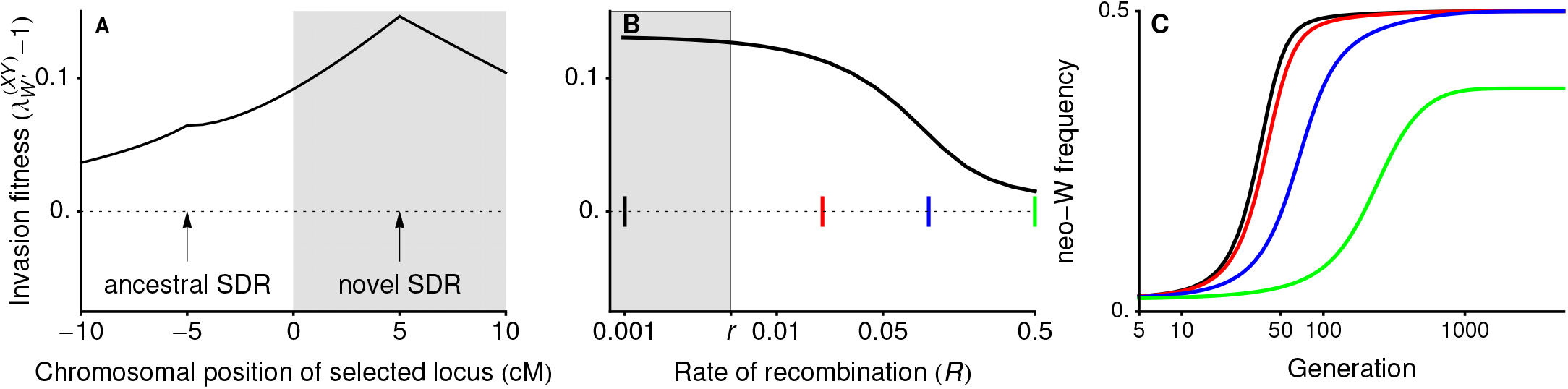
When there is sexually-antagonistic selection and haploid selection, a neo-W allele may invade for any *R*. Panel A shows that the invasion fitness of a neo-W is positive, even when *r* < *R* (unshaded region). In panel B, we vary the recombination rate between the neo-W and the selected locus (*R*) for a fixed recombination rate between the ancestral sex-determining locus and the selected locus (*r* = 0.005). Coloured markers show recombination rates for which the temporal dynamics of neo-W invasion are plotted in panel C (black *R* = 0.001, red *R* = 0.02, blue *R* = 0.1, green *R* = 0.5). The diploid selection parameters used in this plot are the same as in Fig 3. There is also meiotic drive in males favouring 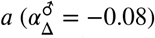, this full set of parameters is marked by an asterisk in Fig S4A. When *R* = 0.5 (green curve), the neo-W does not reach fixation and X, Y, Z, and W alleles are all maintained in the population, see Fig S9C.

**Fig S4.**
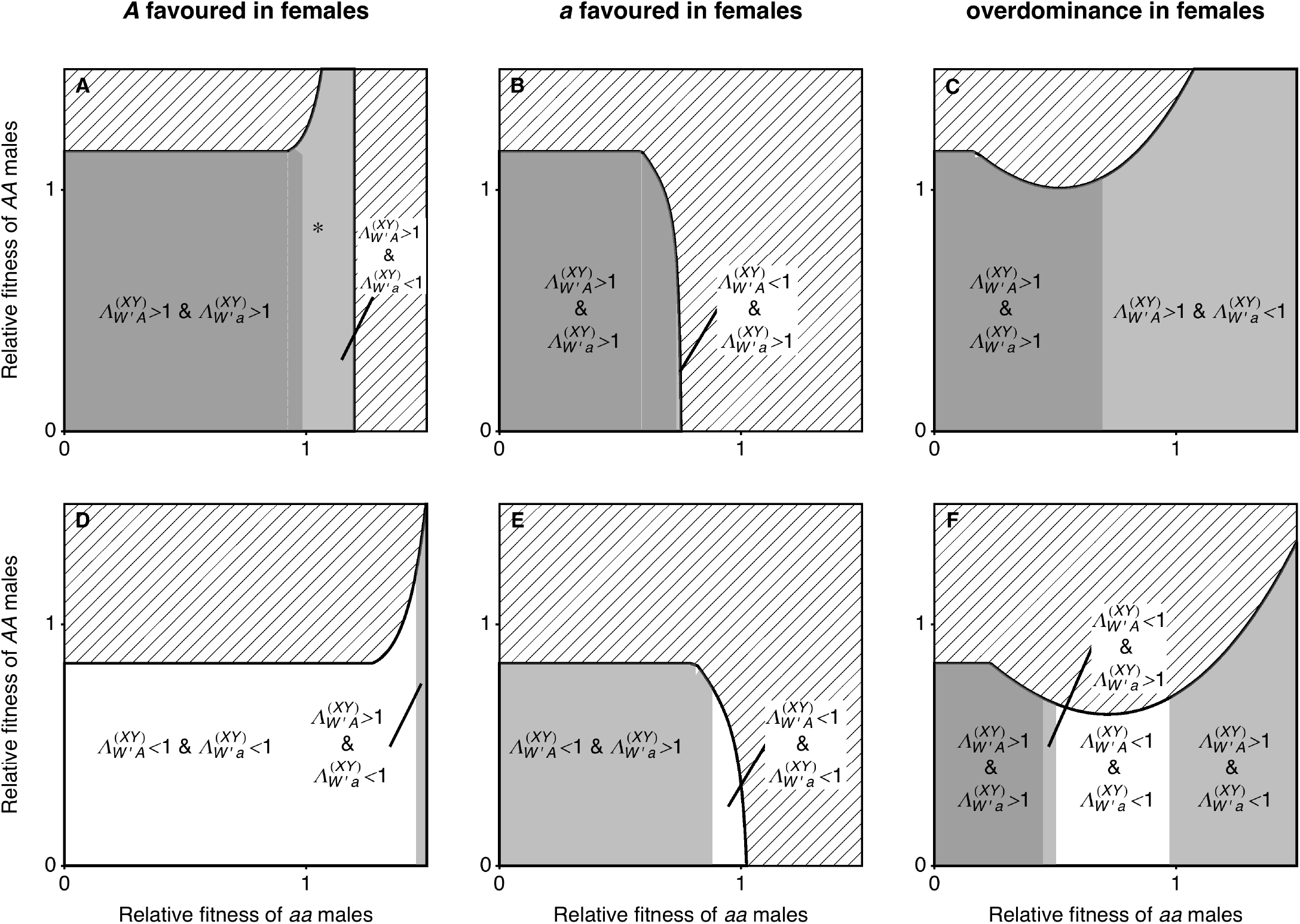
Parameters for which neo-W-*A* and neo-W-*a* haplotypes spread when there is male meiotic drive at a locus that is tightly linked to the ancestral XY locus (*r* = 0). This figure is equivalent to Fig 2 but with meiotic drive in males. In panels A-C, meiotic drive in males favours the *a* allele 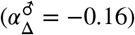, creating male-biased sex ratios and generally increasing 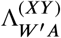 and 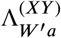. By contrast, 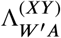 and 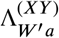 tend to be reduced when meiotic drive in males favours the *A* allele 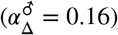, panels D-F.

**Fig S5.**
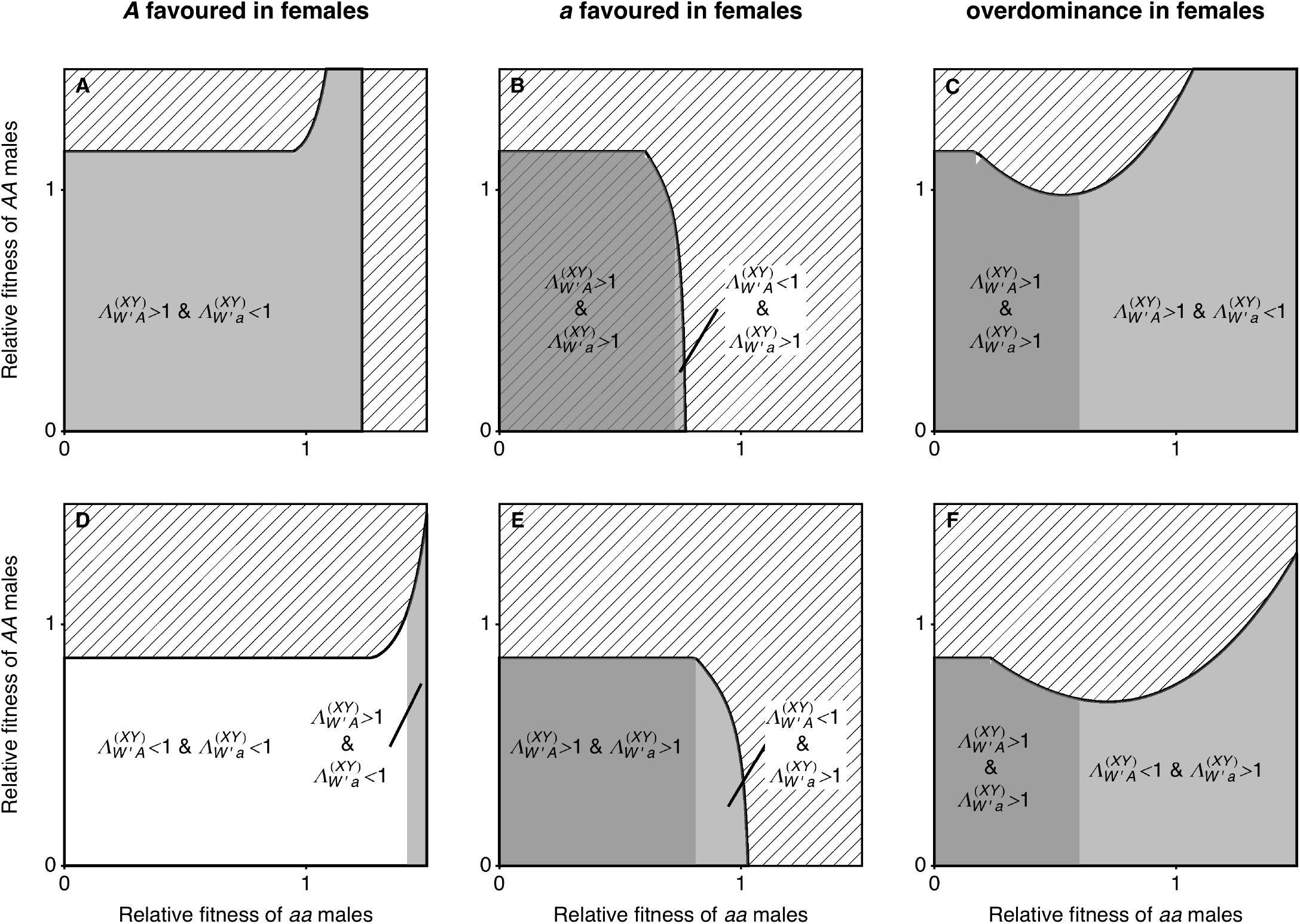
Parameters for which neo-W-*A* and neo-W-*a* haplotypes spread when there is male gametic competition at a locus that is tightly linked to the ancestral XY locus (*r* = 0). This figure is equivalent to Fig 2 but with gametic competition in males. The *a* allele is favoured during male gametic competition in Panels A-C 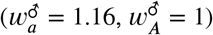, which creates male biased sex ratios and increases 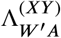 and 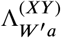. By contrast, 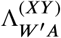 and 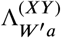 tend to be reduced when the *A* allele is favoured during male gametic competition, panels D-F. Compared to the meiotic drive parameters in Fig S4, the effect of these male gametic competition parameters on the sex ratio is smaller. For example, in Fig S4A-C, the ancestral sex ratio is *α*^♂^ = 0.58 at equilibrium (B) and in panels A-C of this plot, the ancestral sex ratio is 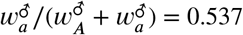 at equilibrium (B).

**Fig S6.**
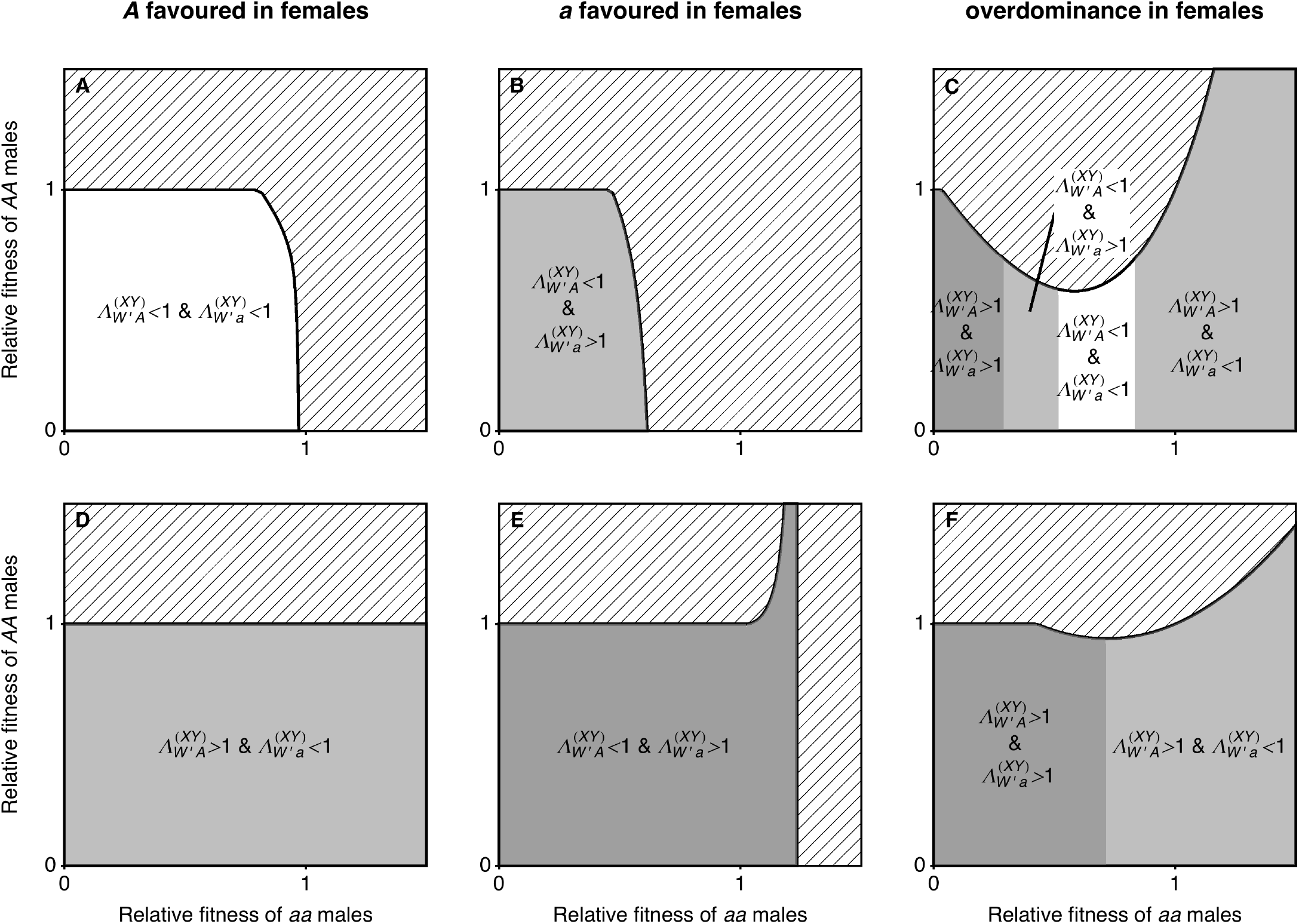
Parameters for which neo-W-*A* and neo-W-*a* haplotypes spread when there is female meiotic drive at a locus that is tightly linked to the ancestral XY locus (*r* = 0). This figure is equivalent to Fig 2 but with meiotic drive in females. The *a* allele is favoured by meiotic drive in females in Panels A-C 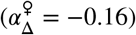, which increases 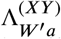 and decreases 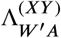 Female meiotic drive in favour of the *A* allele (panels D-F, 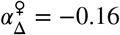) has the opposite effect.

**Fig S7.**
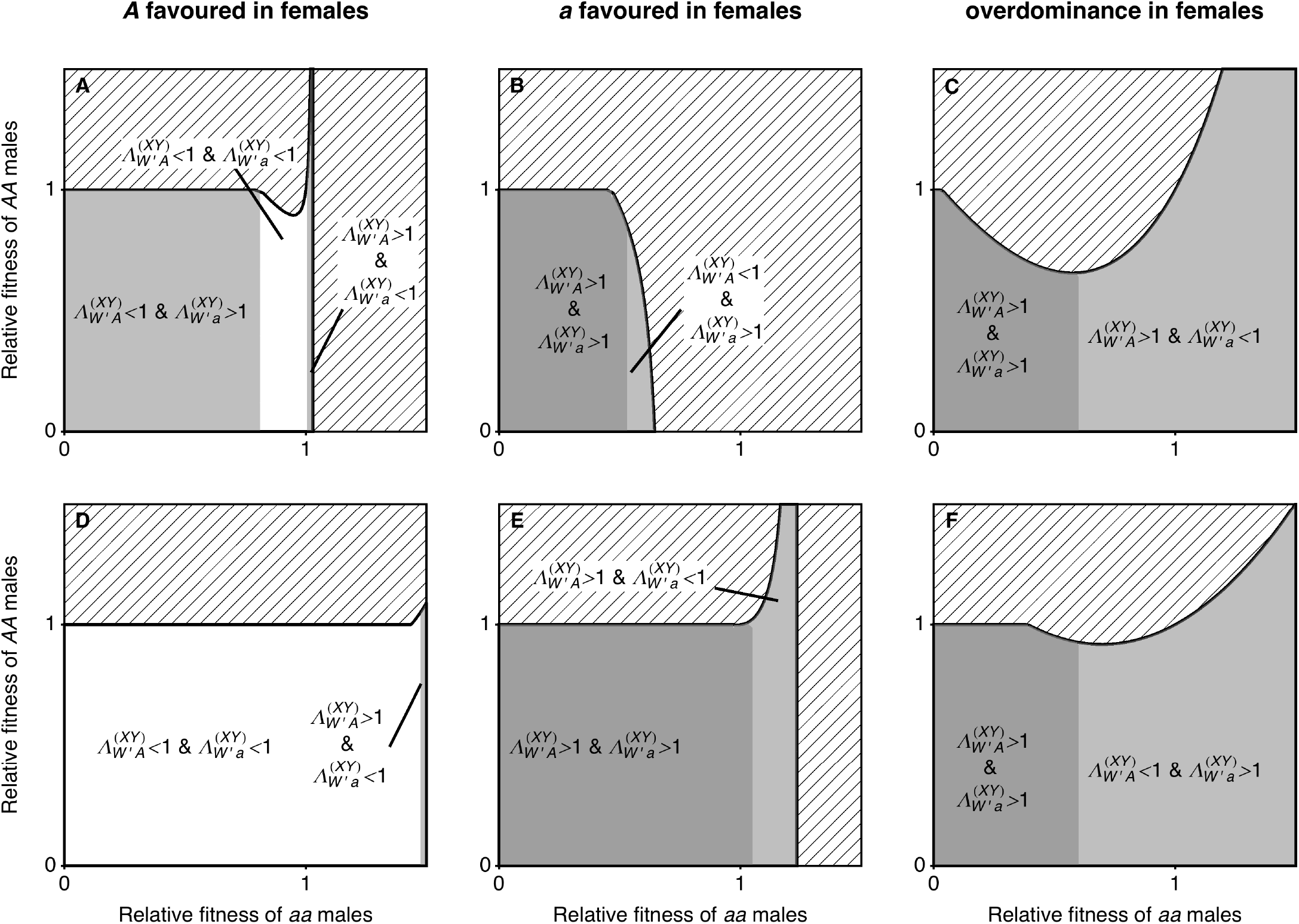
Parameters for which neo-W-*A* and neo-W-*a* haplotypes spread when there is female gametic competition at a locus that is tightly linked to the ancestral XY locus (*r* = 0). This figure is equivalent to Fig 2 but with gametic competition in females. The *a* allele is favoured during female gametic competition in females in Panels A-C 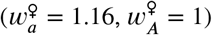, which increases 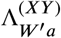 and decreases 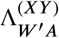. The *A* allele is favoured during gametic competition in panels D-F 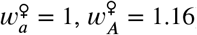, giving the opposite effect on 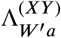 and 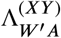.

**Fig S8.**
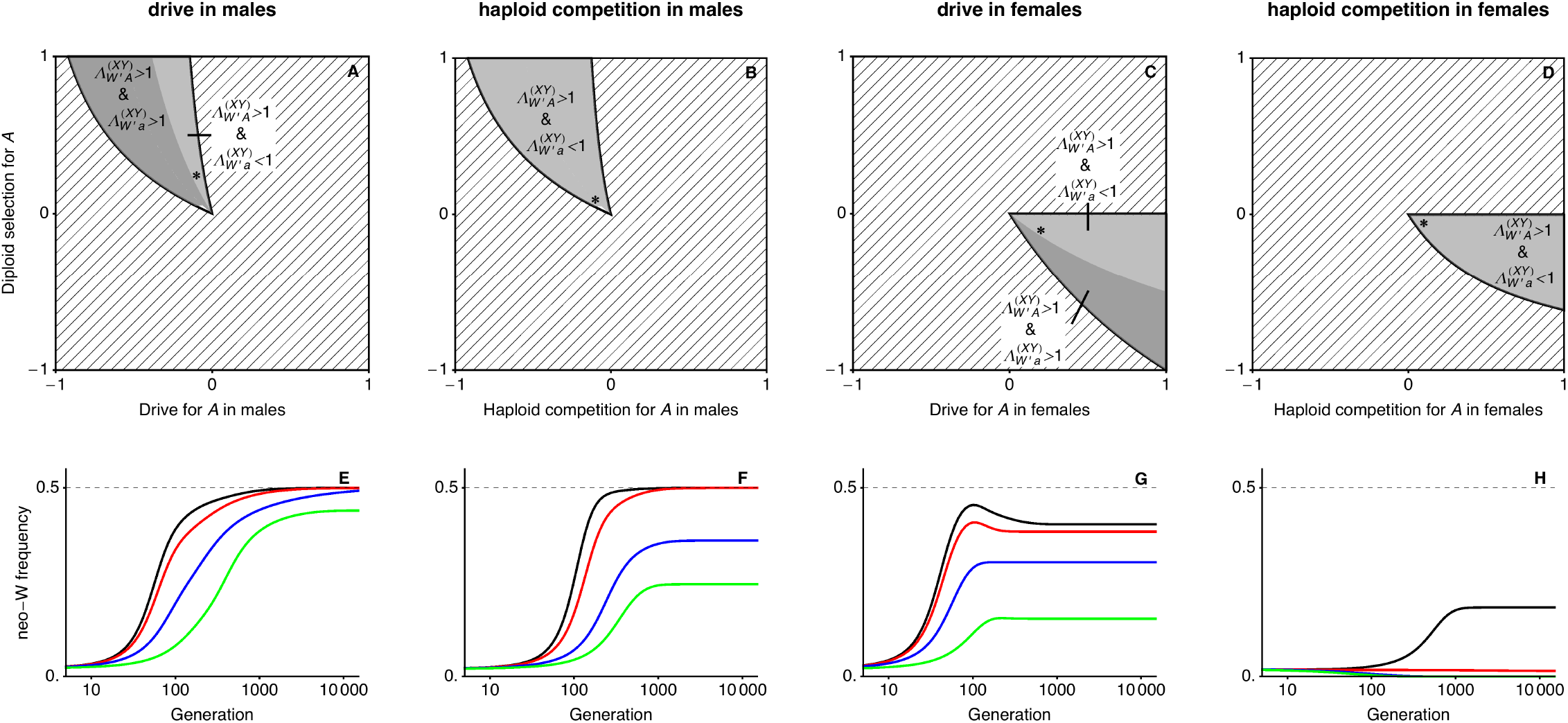
Ploidally-antagonistic selection can drive the spread of neo-W alleles. A-D show when each of the neo-W haplotypes invades an internally stable equilibrium with *a* fixed on the Y (found by setting *r* = 0). The y-axis shows directional selection in diploids of both sexes, *s*^♀^ = *s*^♂^, and the x-axes show sex-limited drive, 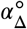, or haploid competition, *t*^∘^. The top left and bottom right quadrants therefore imply ploidally-antagonistic selection (and these are the only places where neo-W haplotypes can invade). Dominance is equal in both sexes, *h*^♀^ = *h*^♂^ = 3/4. E-F show the temporal dynamics of neo-W frequency in females with parameters given by the asterisks in the corresponding A-D plot, with *r* = 1/200, for four different *R*. Black *R* = 1/1000, Red *R* = 2/100, Blue *R* = 1/10, Green *R* = 1/2.

**Fig S9.**
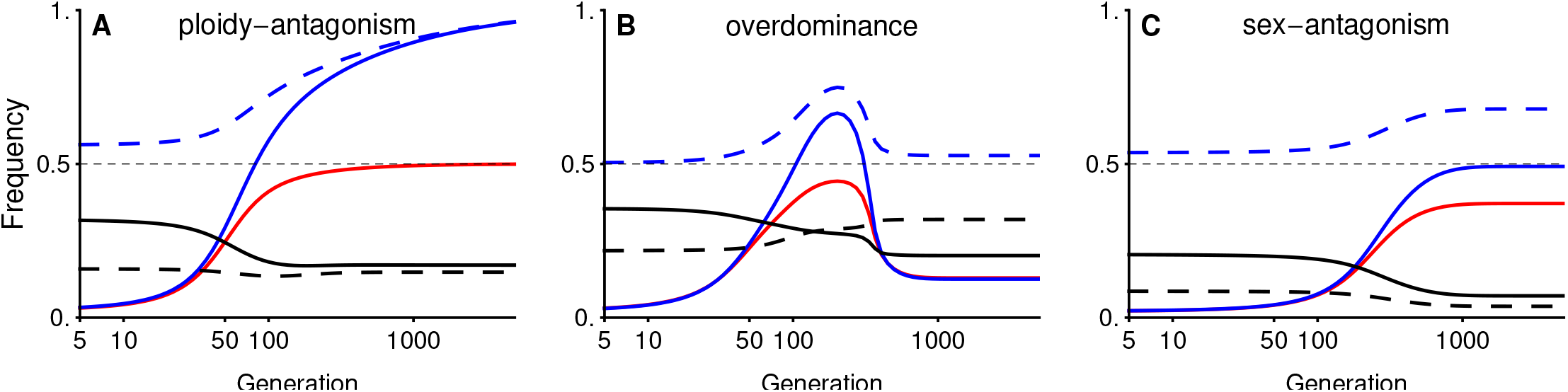
Pseudo-fixation of neo-W or maintenance of multiple sex-determining alleles. The curves show the frequencies of the neo-W (red), ancestral Y (blue), and *A* allele (black) among female gametes (solid curves) and among male gametes (dashed curves). In panel A, there is a complete transition from XY sex determination (XX-ZZ females and XY-ZZ males) to ZW sex determination (YY-ZW females and YY-ZZ males). In panels B and C a polymorphism is maintained at both the ancestral XY locus and the new ZW locus, such that there are males with genotypes XY-ZZ and YY-ZZ and females with genotypes XX-ZZ, XX-ZW, XY-ZW, and YY-ZW. In panel A, selection is ploidally-antagonistic with drive in males (parameters as in the green curve in Fig 5B). In panel B, there is overdominance in both sexes and no haploid selection (parameters as in the green curve in Fig S2C). In panel C, there is sexually-antagonistic selection in diploids with drive in males (parameters as in the green curve in Fig S4C). In all cases, the initial equilibrium frequency has *a* near fixation on the Y.

